# Phosphorylation of the GARP Subunit Vps53 by Snf1 Leads to the Formation of a Golgi – Mitochondria Contact Site (GoMiCS) in Yeast

**DOI:** 10.1101/2020.06.26.173864

**Authors:** Susanne A. Wycislo, Celine Sundag, Stefan Walter, Sebastian Schuck, Florian Fröhlich

## Abstract

The canonical function of the Golgi-associated retrograde protein (GARP) complex is the tethering of transport carriers. GARP belongs to the complexes associated with tethering containing helical rods (CATCHR) family and is a hetero-tetrameric complex consisting of the subunits Vps51, Vps52, Vps53 and Vps54. How the activity of GARP is regulated and if it possesses other functions besides tethering remains largely unknown. Here we identify the GARP subunit Vps53 as a novel regulatory target of the *S. cerevisiae* AMP kinase (AMPK) homolog Snf1. We find that Vps53 is both an *in vivo* and *in vitro* target of Snf1 and show that phosphorylation depends on the nature and quantity of the available carbon source. Phosphorylation of Vps53 does not affect the canonical trafficking pathway, but results in altered mitochondrial dynamics and the formation of a previously unknown contact site between the Golgi apparatus and mitochondria, termed GoMiCS. Our results provide an example of a subunit of a CATCHR complex with a constitutive function in membrane trafficking and an inducible role in organelle contact site formation. We anticipate our results to be the starting point for the characterization of this novel contact site.

## Introduction

Eukaryotic cells are highly compartmentalized into membrane-surrounded organelles to create optimized environments tailored to specialized biochemical reactions. To maintain the specific identity of each organelle, proteins and lipids have to be transported within cells to their correct destination. This complex task is achieved by vesicular trafficking between organelles as well as by a network of membrane contact sites (Bonifacino and Glick, 2004; Guo *et al*., 2014; Eisenberg-Bord *et al*., 2016; Gatta and Levine, 2017). Both vesicular trafficking and the formation of membrane contact sites are highly regulated by the metabolic state of cells (Jones *et al*., 2012; Aoh *et al*., 2013; Hönscher *et al*., 2014; Prinz, 2014).

Two major trafficking pathways operate in cells to maintain the balance between anabolic and catabolic processes, the secretory and the endocytic pathway (Maxfield and McGraw, 2004; Guo *et al*., 2014; Kim and Gadila, 2016). The Golgi apparatus is the central sorting station at the intersection of both pathways. The secretory pathway starts with the delivery of newly synthesized proteins and lipids from the endoplasmic reticulum (ER) to the Golgi apparatus (Dancourt and Barlowe, 2010; Lord *et al*., 2013). Golgi-resident proteins are retained there, while secretory cargo proteins are delivered to the plasma membrane and lysosomal enzymes are sorted towards the endosomal system (De Matteis and Luini, 2008; Guo *et al*., 2014; Kim and Gadila, 2016). In contrast, endocytosis starts with the uptake of proteins and lipids from the plasma membrane which are then sorted in endosomes for recycling or degradation in lysosomes (Pelham, 2002; Doherty and McMahon, 2009). The endosomal recycling pathway leads to the delivery of vesicles from endosomes to the Golgi apparatus (Lewis *et al*., 2000; MacDonald and Piper, 2017). In addition, a direct recycling pathway from the plasma membrane to the Golgi apparatus has been suggested in *S. cerevisiae* (Day *et al*., 2018; Eising *et al*., 2019).

In yeast the Golgi-associated retrograde protein trafficking (GARP) complex has been identified as an important factor for the transport of cargo from endosomes to the Golgi apparatus and potentially from the plasma membrane to the Golgi apparatus (Conibear and Stevens, 2000; Eising *et al*., 2019). GARP is a hetero-tetrameric complex consisting of the four subunits Vps51, Vps52, Vps53 and Vps54 (Conibear and Stevens, 2000, 2003). Structurally GARP belongs to the family of complexes associated with tethering containing helical rods (CATCHR; Vasan *et al*., 2010; Chou *et al*., 2016). This multi-subunit family of protein complexes is associated with tethering of endosomal transport carriers. The GARP complex is recruited to the Golgi apparatus by the Rab GTPase Ypt6 and interacts with the SNARE (soluble N-ethylmaleimide-sensitive-factor attachment receptor) protein Tgl1 via its Vps51 subunit (Siniossoglou and Pelham, 2001; Conibear and Stevens, 2003). The GARP complex has been linked to a multitude of intracellular processes. Besides its canonical role in sorting the carboxypeptidase Y (CPY) receptor Vps10, the GARP complex is important for sphingolipid homeostasis, autophagy, mitochondrial tubulation, and vacuolar integrity (Conibear and Stevens, 2000; Reggiori and Klionsky, 2006; Fröhlich *et al*., 2015; Yang and Rosenwald, 2016).

Regardless of its involvement in many cellular processes, regulation of the GARP complex or of its subunits by post-translational modifications has not been described. However, previous studies have identified different phosphorylation sites on subunits of the GARP complex (Gnad *et al*., 2009; Braun *et al*., 2014; Fröhlich *et al*., 2016). Based on results showing differential phosphorylation after sphingolipid depletion, we decided to characterize a phosphorylation site in the carboxy-terminus of the GARP subunit Vps53.

Here, we use a combination of sequence prediction, mass spectrometry-based proteomics, and *in vitro* kinase assays to identify the yeast AMP kinase homolog Snf1 as a regulator of Vps53. We demonstrate that Vps53 phosphorylation does not affect the canonical endosomal recycling pathway mediated by GARP. Using mass spectrometry-based proteomics and live cell imaging, we describe an unexpected role for Vps53 phosphorylation in mitochondrial dynamics which is mediated by the formation of a previously unknown membrane contact site formed between the Golgi apparatus and mitochondria.

## Results

### Vps53 is phosphorylated by Snf1 at position 790 *in vivo*

We and others have previously identified phosphorylated residues on subunits of the GARP complex (Gnad *et al*., 2009; Braun *et al*., 2014; Fröhlich *et al*., 2016). Given the essential nature of GARP in endo-lysosomal trafficking and sphingolipid homeostasis, we decided to test if phosphorylation is a potential regulatory mechanism. Especially the Vps53 subunit has been shown multiple times to be phosphorylated at serine residue 790. This residue is located at the very c-terminus of Vps53 which is functionally accessible to kinases and is important for the function of Vps53 (Vasan *et al*., 2010; Chou *et al*., 2016).

Therefore, we first analyzed the amino acid sequence surrounding serine 790 in the Vps53 subunit of GARP. Carrying an arginine residue in the −3 position as well as hydrophobic residues in the −10, −5 and +4 position, it strongly resembles a classical motif for AMP activated protein kinase (AMPK) or its homolog Snf1 in *S. cerevisiae* (Dale *et al*., 1995; Hardie, 2007) (Fig. 1a). In yeast, Snf1 is not activated by adenosine monophosphate (AMP) levels but is repressed by glucose (Jiang and Carlson, 1996; Wilson *et al*., 1996). Depletion of glucose from the growth medium therefore results in increased Snf1 activity. We used stable isotope labeling by amino acids in cell culture (SILAC; Ong *et al*., 2002) combined with affinity purification and mass spectrometry based proteomics to determine if Vps53 phosphorylation was dependent on Snf1 (Fig. 1b). Affinity purification of Vps53 from cells labeled with lysine (K0), grown in 2% glucose compared to ^13^C_6_^15^N_2_-lysine (K8) labeled cells grown for 15 minutes in the presence of 0.02% glucose resulted in the identification of the three GARP subunits Vps52, Vps53 and Vps54. The heavy/light ratio of the three purified proteins was close to 1, suggesting that glucose depletion in the medium neither changed the abundance of the proteins in the cell nor their assembly into the GARP complex (Fig. 1c). In contrast, the peptide of Vps53 carrying a phosphorylated serine at position 790 (S790) showed a high heavy/light ratio, while the corresponding non-phosphorylated peptide showed a ratio smaller than 1 (Fig. 1c). The acquired data even allowed us to calculate that Vps53 phosphorylation occupancy at serine 790 increased from ∼ 10% in glucose grown cells to approximately 45% in glucose depleted cells (Fig. 1d). Together, these results indicate that Vps53 is phosphorylated upon glucose deprivation.

**Figure 1:**
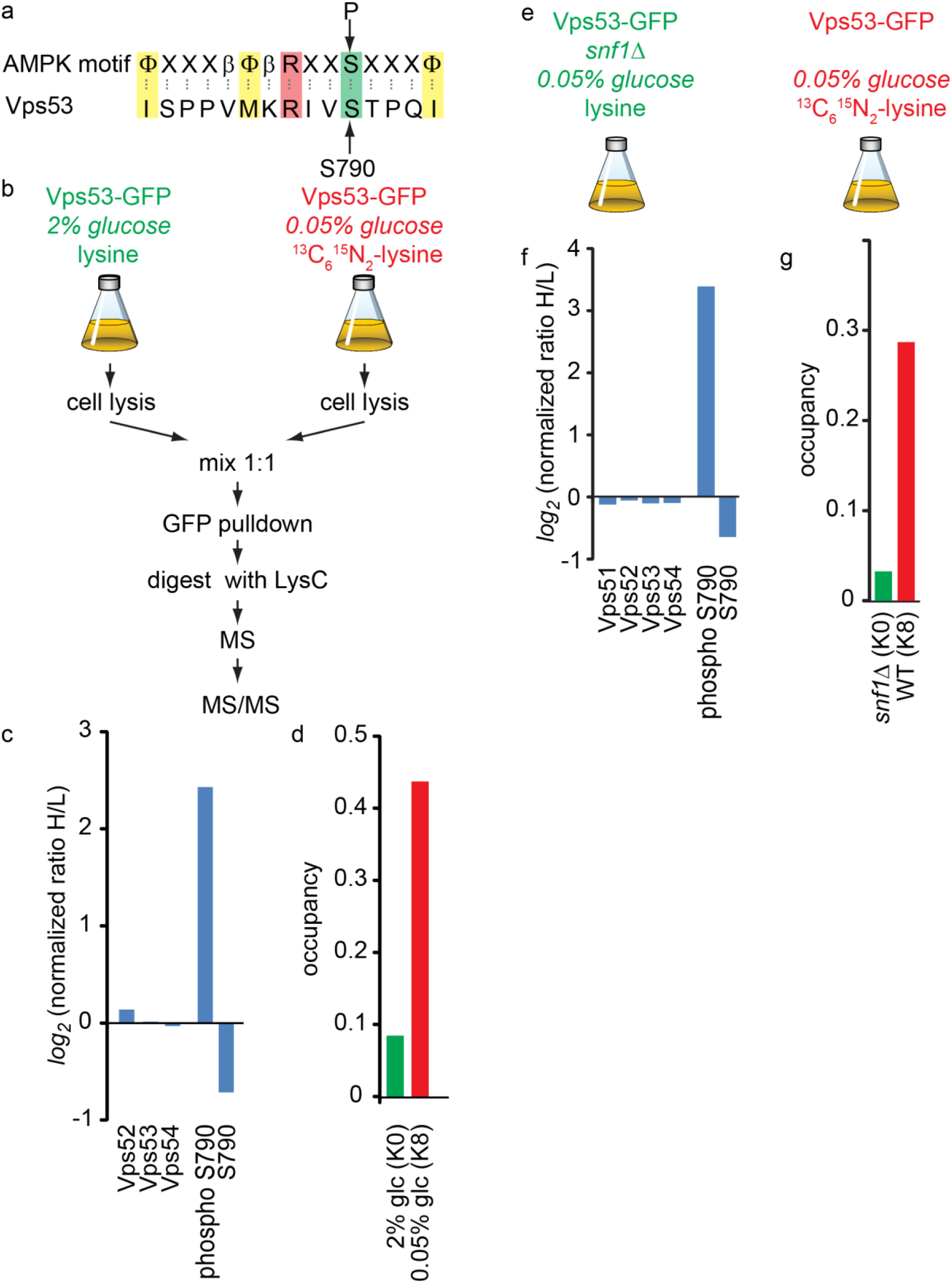
Vps53 is phosphorylated by AMPK *in vivo*. **(a)** Vps53 has a c-terminal has a c-terminal AMPK phosphorylation motif. The AMPK consensus motif has a sequence of β φ βXXXS/TXXXφ (basic, β = R, K or H; hydrophobic, φ = M, L, I, F or V; X = any amino acid; S/T = phosphorylation site), **(b)** Mass spectro-metric SILAC setup for *in vivo* Vps53 pulldowns. **(c**,**d)** Quantification of Vps53 phosphorylation from heavy labeled glucose starved cells and light labeled control cells, **(c)** Vps53-GFP purified from SILAC-labeled cells grown with or without glucose was analyzed by MS. log2 transformaed normal ratios for proteins Vps52, Vps53 and Vps54 are shown. Unphosphorylated Vps53 peptide (S790) and its phosphorylated form (phospho S790) are shown **(d)** Quantification of Vps53 phosphorylation occupancy at serine 790 is shown. Phosphorylation occupancy in glucose grown cells (green) and glucose starved cells (red) is plotted, **(e)** Experimental setup to determine Snf1 dependent Vps53 phosphorylation. Vps53-GFP purified from SILAC-labeled *snf1Δ* (light) or WT cells (heavy) grown without glucose was analyzed by MS. **(f)** Vps53-GFP purified from SILAC-labeled snf1D or WT cells starved for glucose was analyzed by MS. log2 transformaed normal ratios for proteins Vps51, Vps52, Vps53 and Vps54 are shown. Unphosphorylated Vps53 peptide (S790) and its phosphorylated form (phospho S790) are Shown **(g)** Quantification of Vps53 phosphorylation occupancy at serine 790 is shown. Phosphorylation occupancy in *snf1Δ* cells (green) and WT cells (red) is plotted.

To test if this phosphorylation is indeed depending on Snf1, we affinity purified Vps53-GFP from lysine labelled *snf1Δ* cells and from ^13^C_6_^15^N_2_-lysine labelled WT cells, both grown for 15 min in the presence of 0.02% glucose (Fig. 1e). In these experiments we were able to co-purify all four subunits of the GARP complex, Vps51, Vps52, Vps53 and Vps54. None of the above showed a difference in total protein levels, suggesting that the abundance of the proteins as well as the formation of the complex does not depend on Snf1 (Fig. 1f). However, we observed a high heavy/light ratio for the phosphorylated Vps53 peptide including serine 790 with a concomitant low ratio of the non-phosphorylated peptide (Fig. 1f). We calculated the amount of phosphorylated Vps53 in *snf1Δ* cells to be 2% (which is likely an overestimate, see material and methods) and approximately 30% in WT cells (Fig. 1 e).

In a previous study we have identified a decrease in Vps53 phosphorylation at position 790 upon chemical inhibition of sphingolipid biosynthesis with myriocin (Fröhlich *et al*., 2016). Our results indicate that Vps53 phosphorylation is very low under standard growth conditions. Therefore, a myriocin dependent reduction probably reflects minor changes. However, we tested Snf1 phosphorylation upon myriocin treatment, glucose starvation and a combination of both treatments by western blotting. As expected, myriocin dependent sphingolipid depletion had only minor effects on basal Snf1 activity based on phosphorylation of threonine residue 210. A combination of myriocin treatment and glucose starvation led to a delayed decrease of Snf1 activity (suppl. Fig. 1).

Together, our data suggest that Vps53 is a target of the yeast AMPK homolog Snf1 and is phosphorylated upon depletion of the carbon source in the growth medium. Sphingolipid depletion only shows minor effects on basal Snf1 activity.

### Vps53 is a substrate of Snf1 *in vitro*

To determine if Vps53 is a substrate of Snf1 *in vitro*, we first purified the C-terminal part of Vps53 (amino acids 552-882) recombinantly expressed in *E*.*coli* (Fig. 2a). In addition, we purified the recombinantly-expressed kinase domain of Snf1 (aa 1-392) as well as a non-activatable Snf1 catalytic domain carrying an alanine residue at position 210 instead of a threonine (Fig. 2b). Snf1 is activated by phosphorylation on threonine 210 by either Sak1, Tos3, or Elm1 (Hong *et al*., 2003). To activate Snf1 *in vitro*, we purified TAP-tagged Elm1 lacking the autoinhibitory c-terminal domain (residues 421-640) from yeast cells. Incubation of Snf1 and purified Elm1 results in the phosphorylation of Snf1 at position 210 (Lee *et al*., 2012). While *in vitro* kinase assays are usually performed in the presence of radio-labeled ATP, we decided to develop a label free mass spectrometry-based Snf1 *in vitro* kinase assay with site-specific accuracy. For this purpose, we incubated the purified C-terminus of Vps53 with either the Elm1-activated Snf1_WT_ or kinase-dead (KD) Snf1_T210A_ in triplicates and analyzed the resulting peptides after digestion with the endo-proteinase trypsin by mass spectrometry (Fig. 2c). This analysis resulted in the complete identification of all peptides from the C-terminus of Vps53 as well as the entire kinase domain of Snf1. We also detected multiple phosphorylated residues on Vps53. However, we only identified the Vps53 peptide carrying the phosphorylated amino acid residue serine 790 (RIVpSTPQIQQQK) in the presence of the activated kinase domain of Snf1 (Fig. 2d). Other phosphorylation events are therefore probably artifacts resulting from phosphorylation by purified Elm1 or other impurities from this purification. Our experiments clearly demonstrate that Vps53 is a substrate of Snf1, both *in vivo* and *in vitro*.

**Figure 2:**
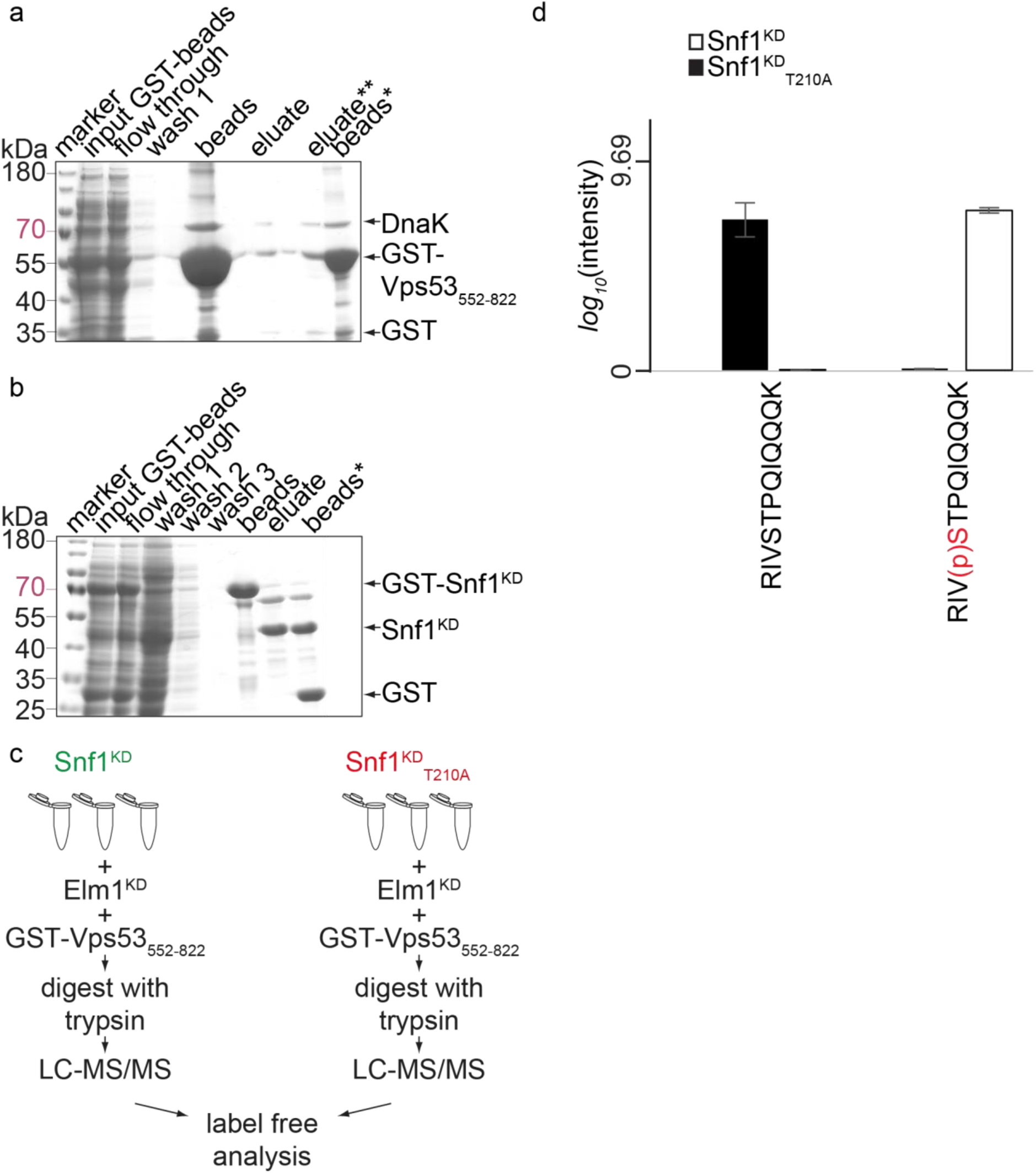
Vps53 is a Snf1 substrate *in vitro*. **(a,b)** Overexpressed recombinant GST tagged *Vps53*_*552822*_ was purified from E.coli. SDS-PAGE showing the lysates and affinity purification steps. Before (beads) and after elution (beads*) of glutathione agarose matrix, 2 % were boiled in Laemmli buffer to elute protein. 0.2 % of the lysate (input GST-beads) and lysate after affinity purification (flow through) were loaded on a 10 % SDS gel. Left lane shows the molecular weight markers (marker), **(a)** Recombinant *Vps53*_*552822*_ was twice eluted via glutathione (eluate (15 mM), eluate*(20 mM)). 2% of the eluates were loaded. **(b)** GST-Snfl kinase domain was eluted following PreScission protease digestion. eluate: eluted Snf1-KD **(c)** Experimental workflow of label free, mass spectrometry based *in vitro* kinase assay to determine kinase-substrate specificity of Snf1 and Vps53. **(d)** Vps53 is a substrate of Snf1 *in vitro*. Average peptide intensities of the Vps53 peptide RIVSTPQIQQQK (left) and its phosphorylated counterpart (right) from experiments with the WT Snf1 kinase domain (white bars) or the inactive Snf1 kinase domain (black bars). Error bars represent standard deviation from three experiments.

### Phosphorylation of Vps53 is functional for GARP complex assembly and localization

Next, we asked which physiological function of Vps53 or the GARP complex is affected by phosphorylation of Vps53. Our analysis of the purified GARP complex showed that the assembly of the complex was unaffected by glucose limiting conditions and concomitant phosphorylation of Vps53 (Fig. 1d). To confirm these results, we generated GFP-tagged versions of Vps53, where serine 790 is exchanged to either alanine (S790A, non-phosphorylatable) or to aspartate (S790D, phospho-mimetic). We performed affinity purifications of lysine 8 labelled Vps53-GFP, *Vps53*_*S790A*_-GFP and *Vps53*_*S790D*_-GFP cells compared to lysine 0 labelled WT cells (Fig. 3a). Under the used conditions we were able to co-purify the three GARP subunits Vps52, Vps53 and Vps54 in cells expressing Vps53-GFP, *Vps53*_*S790A*_-GFP and *Vps53*_*S790D*_-GFP, confirming that the assembly of the GARP complex is unaffected by Snf1 dependent phosphorylation on serine 790 (Fig. 3b, c and d). To further confirm these results, we analyzed the localization of mKate tagged Vps53, *Vps53*_*S790A*_, and *Vps53*_*S790D*_ in cells relative to Vps54 tagged with neonGreen (Fig. 3e). We did not observe changes of the localization of either Vps53 construct in comparison to the GARP subunit Vps54 (Fig. 3e). Together, our results show that phosphorylation of Vps53 does not affect GARP complex assembly or localization.

**Figure 3:**
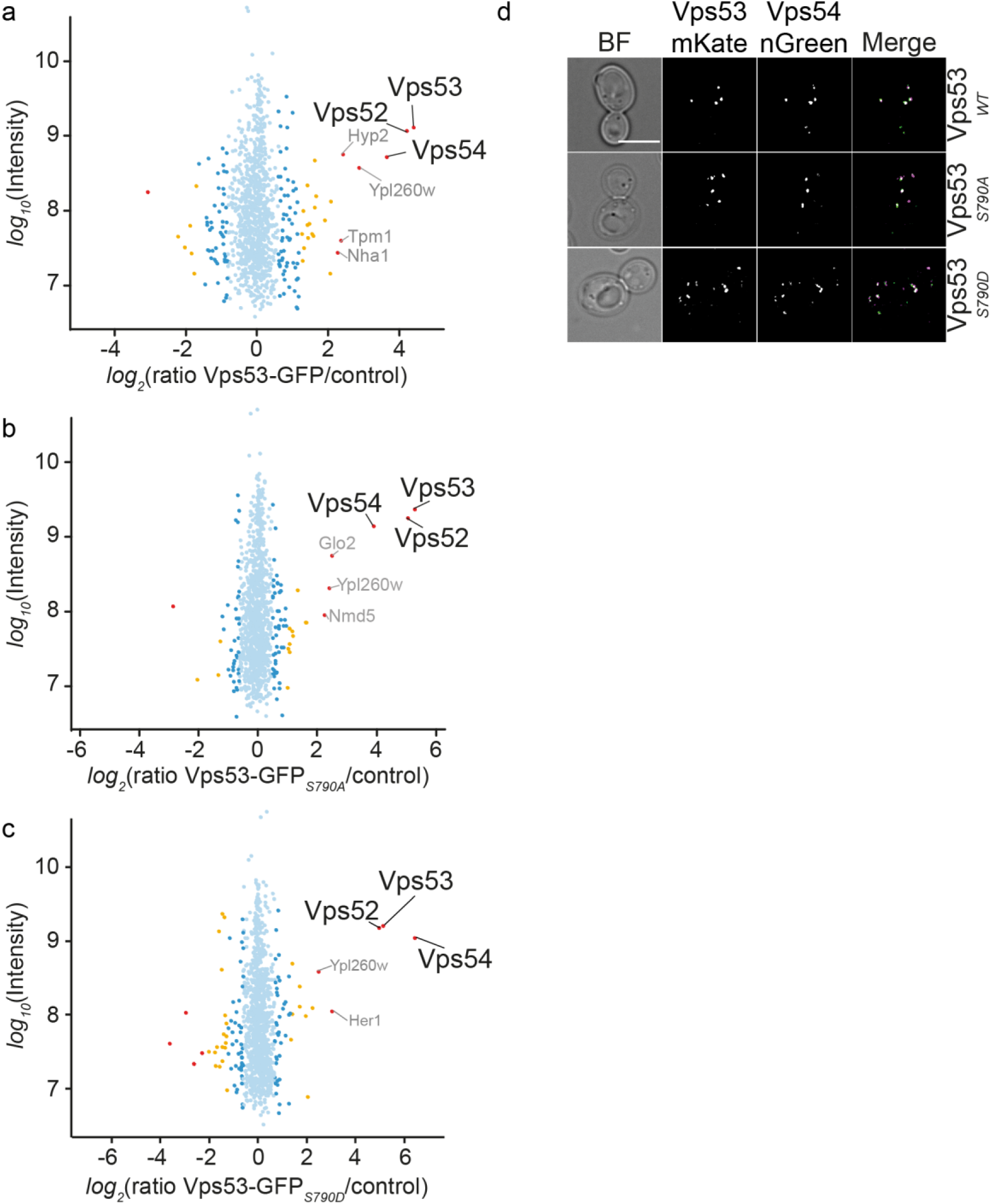
Phosphorylation of Vps53 is functional for GARP complex assembly and localization. **(a)** Affinity purification and MS analysis of “heavy”-labeled” cells expressing GFP-tagged Vps53 and untagged control cells. Intensities are plotted against normalized heavy/light SILAC ratios. Significant outliers (p < 1e-11) are colored in red, orange (p < 0.0001), or blue (p < 0.05); other identified proteins are shown in light blue, **(b)** Affinity purification and MS analysis of “heavy”-labeled” cells expressing GFP-tagged *Vps53*_*S790A*_ and untagged control cells. Intensities are plotted against normalized heavy/light SILAC ratios. Significant outliers (p < 1e-11) are colored in red, orange (p < 0.0001), or blue (p < 0.05); other identified proteins are shown in light blue, **(c)** Affinity purification and MS analysis of “heavy”-labeled” cells expressing GFP-tagged *Vps53*_*S790D*_ and untagged control cells. Intensities are plotted against normalized heavy/light SILAC ratios. Significant outliers (p < 1e-11) are colored in red, orange (p < 0.0001), or blue (p < 0.05); other identified proteins are shown in light blue, **(d)** Vps53 phosphorylation does not affect co-localization with Vps54. Vps53wt (upper panels) *Vps53*_*S790A*_ (middle panels) and *Vps53*_*S790D*_ (lower panels) tagged with mKate (left panels) were co-localized with mNeonGreen tagged Vps54. Merged chanells in the right panels. Sacle bar = 5μM

### Carbon source dependent phosphorylation of Vps53 is functional for the CPY trafficking pathway

Next, we asked if the canonical function of the GARP complex in CPY trafficking is affected by the phosphorylation state of Vps53. Therefore, we used a CPY secretion assay based on a CPY-invertase fusion protein, which allows secretion to be quantified by a colorimetric assay (Robinson *et al*., 1988; Darsow *et al*., 2000; Bean *et al*., 2017). Since this assay detects glucose levels resulting from the hydrolysis of sucrose into fructose and glucose, we had to adjust our growth conditions. For conditions, where Snf1 is inactive and thus Vps53 is non-phosphorylated, we grew cells in the presence of 2% fructose. For Snf1 activating conditions we switched cells to 2% lactate as this had been reported to activate Snf1 (Defenouillère *et al*., 2019). We tested CPY secretion in *vps53Δ* cells expressing a Vps53 WT version, the non-phosphorylatable *Vps53*_*S790A*_ or the phosphomimetic *Vps53*_*S790D*_ mutant as the sole Vps53 copy from a plasmid. We detected a small increase in CPY secretion based on invertase activity in lactate grown cells compared to fructose grown cells. However, we could not observe major differences in the two phospho-mutants compared to WT cells (Fig. 4a). In addition, the levels of CPY secretion were at least ten times lower than in *vps53Δ* cells not expressing a copy of Vps53 (Fig. 4b). Thus, we conclude that the canonical GARP dependent CPY trafficking pathway is not regulated by carbon source and Vps53 phosphorylation by Snf1.

**Figure 4:**
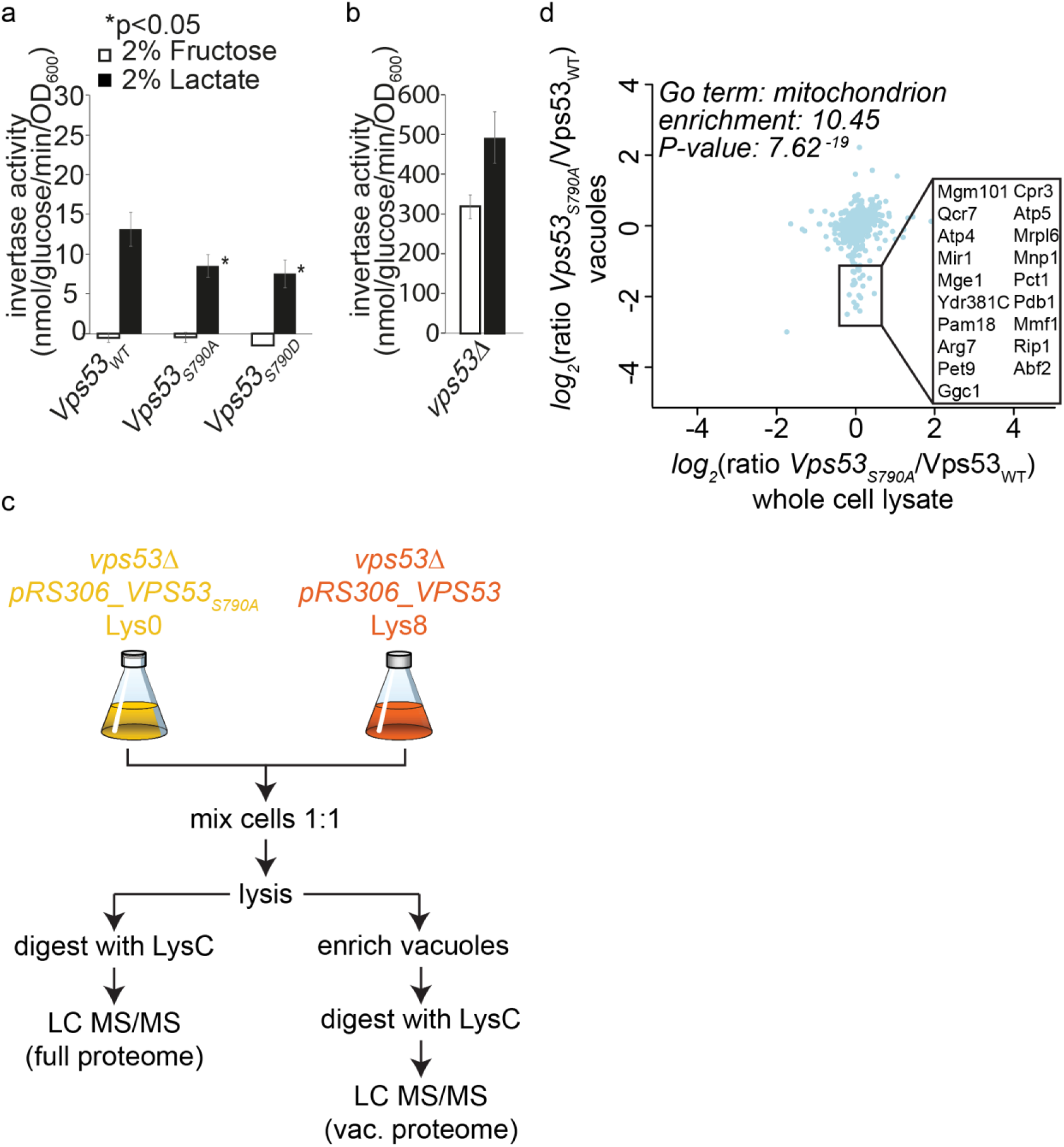
Vps53 phosphorylation does not affect endosomal trafficking. **(a)** *Vps53*_WT_, *Vps53*_*S790A*_ and *Vps53*_*S790D*_ cells showed no CPY secretion in fructose contaiing medium (white bars). Growth in lactate containing media (black bars) resulted in slightly elevated CPY secretion, **(b)** Increaseed levels of secreted invertase activity were detected in vps53Δ cells in fructose grown cells (white bars) and lactate grown cells (black bars), **(c)** Experimental setup to determine proteins mis-sorted to the vacuole in *Vps53*_*S790A*_ cells compared to WT cells grown for 90 min in the presence of lactate. **(d)** Mitochondria are co-enriched in Vps53 purified vacuoles under respiratory growth conditions. SILAC ratios from the cell lysates are plotted against SILAC ratios from the vacuoles of K0 labelled *VPS53*_*S790A*_ cells and K8 lablled *VPS53* cells. Detected mitochondrial protein-levels are decreased in *VPS53*_*S790A*_ purified vacuoles compared to WT. Enriched GO term were identified with the Gene Ontology enRichment anaLysis and visuaLizAtion tool, GOrilla.

In addition to its canonical function in Vps10 recycling and the CPY pathway, we have recently shown that the induced depletion of Vps53, and thus the GARP complex, results in the re-localization of several plasma membrane proteins to the vacuole (Eising *et al*., 2019). We hypothesized that we can identify any cargo whose transport to the vacuole depends on Vps53 phosphorylation using this assay. We therefore purified vacuoles from lysine 8 labelled *VPS53*_*S790A*_ cells and light lysine labelled WT cells and compared their proteomes by mass spectrometry. To account for differences in protein abundance we also analyzed the total proteome of these cells. In addition, we switched cells from glucose containing medium to lactate containing medium for 90 minutes (Fig. 4c). We assumed that only Vps53 from WT cells is phosphorylated under these conditions and also kept conditions comparable to the CPY secretion assay. When we plotted normalized ratios of proteins from the entire cell lysate against the ratios from vacuole isolations we identified a group of proteins that are specifically depleted in vacuole isolations of *VPS53*_*S790A*_ cells (Fig. 4d). We did not identify any cargo from the endo-lysosomal pathway (e.g. Vps10, Prc1, Dnf1, Lem3). In contrast, the group of specifically depleted proteins are almost exclusively mitochondrial proteins, based on GO term enrichments (Fig. 4d and supplementary table 1). The identified mitochondrial proteins were not specific for any mitochondrial compartment (outer membrane, inner membrane, matrix), suggesting that intact mitochondria co-purified with vacuoles in WT cells and that this co-purification is diminished in *VPS53*_*S790A*_ cells (Fig. 4d and supplementary table 1).

**Table 1.**
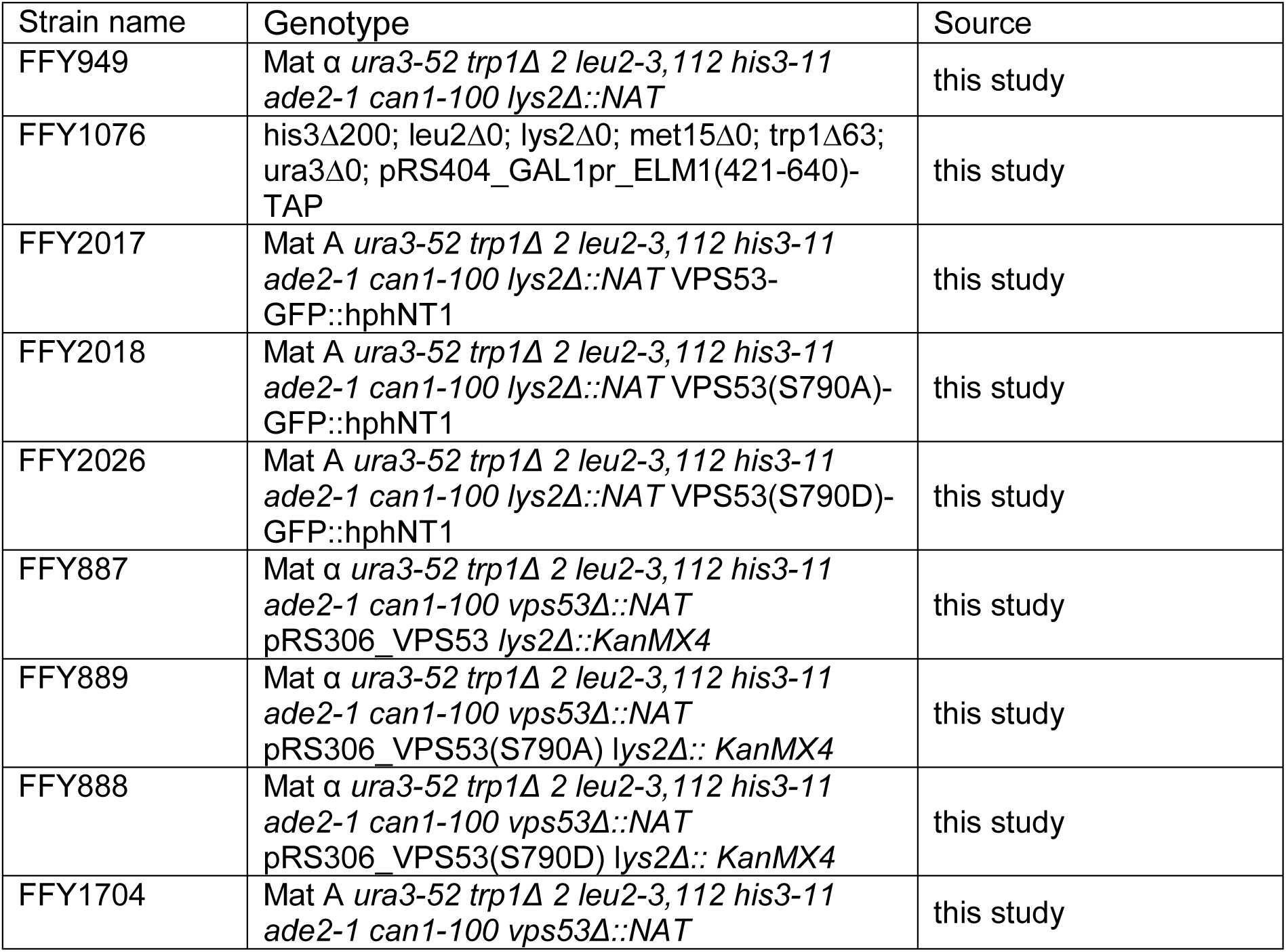
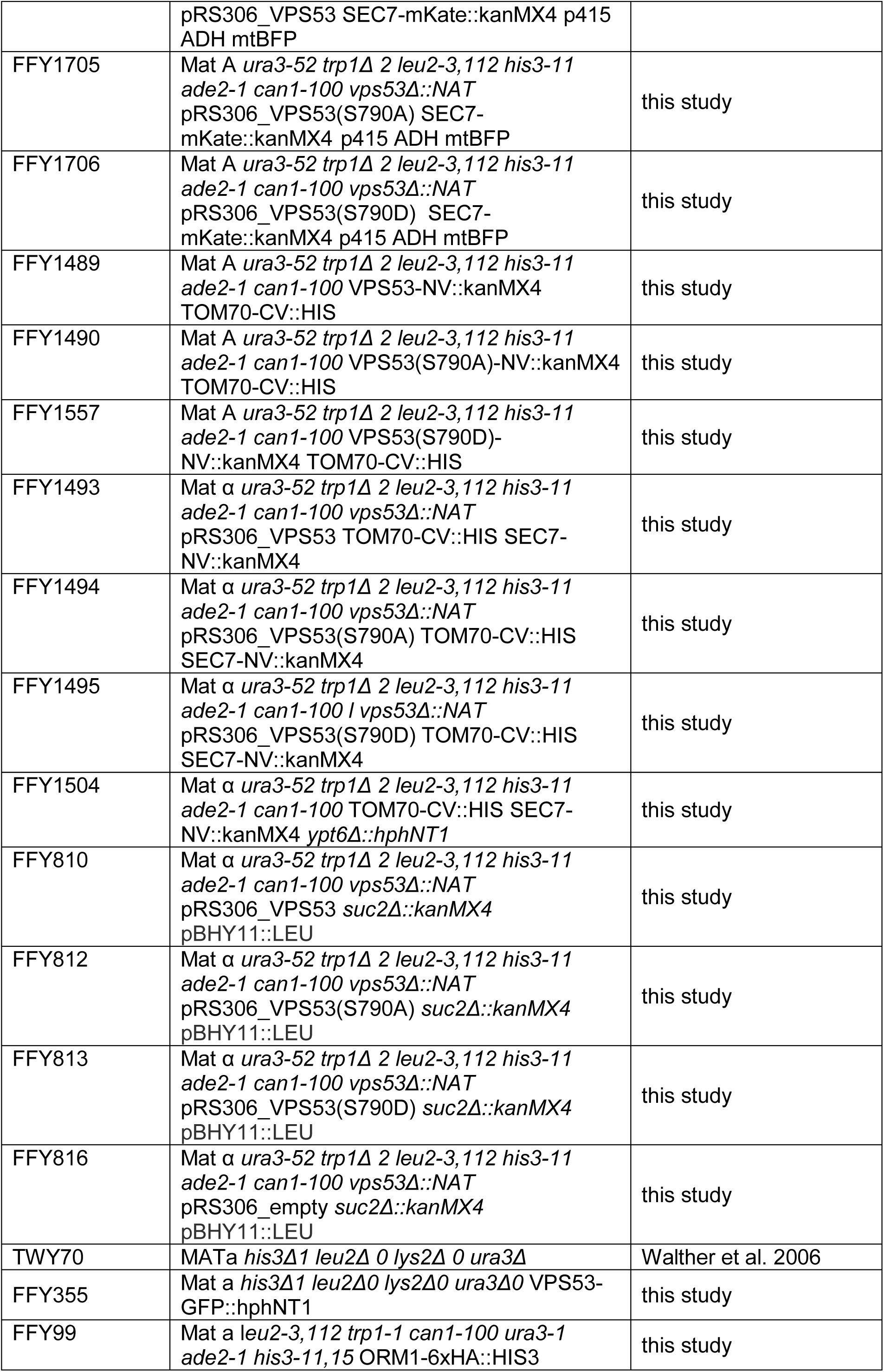
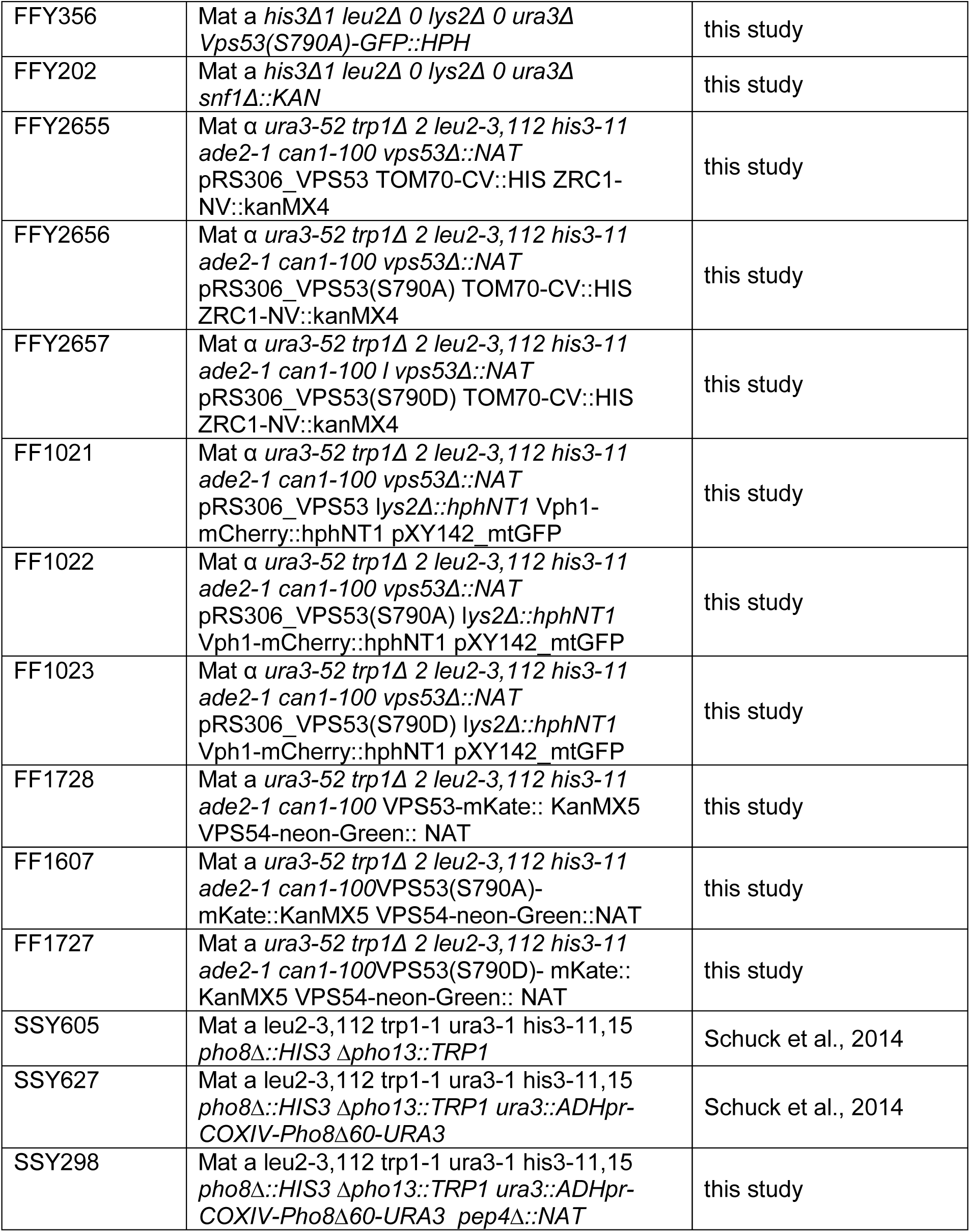
Yeast strains

**Table 2.**
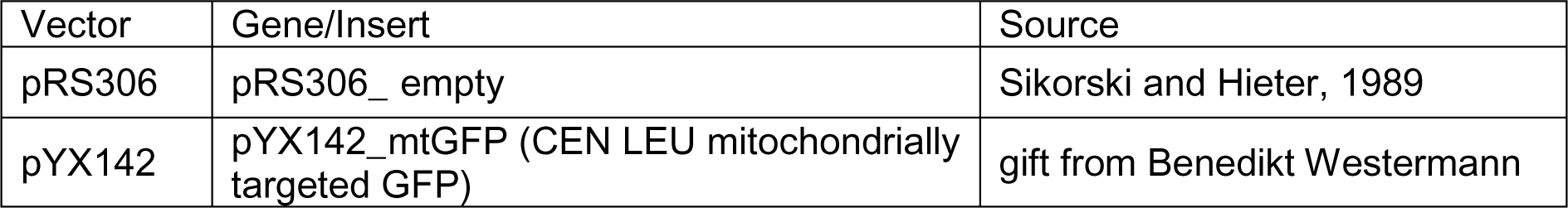
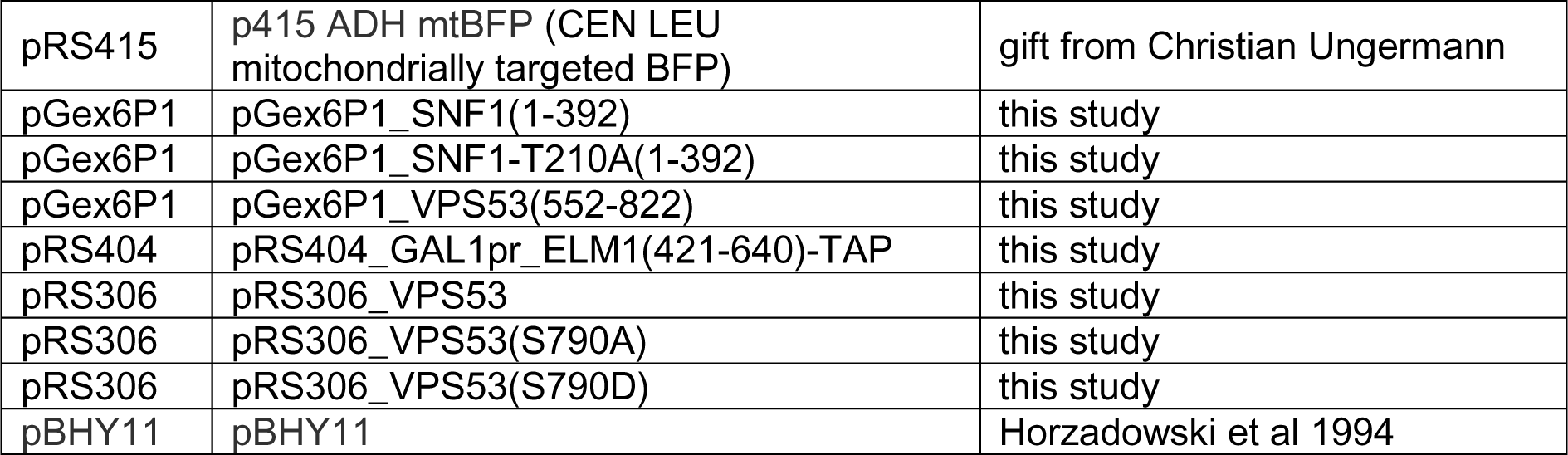
Plasmids.

| Vector | Gene/Insert | Source |
| --- | --- | --- |
| pRS306 | pRS306_ empty | Sikorski and Hieter, 1989 |
| pYX142 | pYX142_mtGFP (CEN LEU mitochondrially targeted GFP) | gift from Benedikt Westermann |
| pRS415 | p415 ADH mtBFP (CEN LEU mitochondrially targeted BFP) | gift from Christian Ungermann |
| pGex6P1 | pGex6P1_SNF1(1-392) | this study |
| pGex6P1 | pGex6P1_SNF1-T210A(1-392) | this study |
| pGex6P1 | pGex6P1_VPS53(552-822) | this study |
| pRS404 | pRS404_GAL1pr_ELM1(421-640)-TAP | this study |
| pRS306 | pRS306_VPS53 | this study |
| pRS306 | pRS306_VPS53(S790A) | this study |
| pRS306 | pRS306_VPS53(S790D) | this study |
| pBHY11 | pBHY11 | Horzadowski et al 1994 |

### Phosphorylation of Vps53 affects mitochondrial dynamics and morphology

To uncover the mechanistic basis of decreased mitochondrial co-purification in *Vps53*_*S790A*_ cells we analyzed mitochondrial morphology under both, fermentative and respiratory growth conditions. First we defined three different mitochondrial phenotypes in cells, tubulated, intermediate or fragmented based on mito-GFP fluorescence (Fig. 5a; Rambold *et al*., 2011; Aung-Htut *et al*., 2013). Similarly to previous studies, about 80% of cells expressing a WT copy of Vps53 had tubulated mitochondria when grown in glucose (Wong *et al*., 2000). This number similar in *Vps53*_*S790D*_ cells and slightly reduced to approximately 60% in *VPS53*_*S790A*_ expressing cells with a concomitant increase in mitochondria showing an intermediate morphology (Fig. 5a, black bars). Switching cells to lactate for 90 min resulted in mitochondrial fragmentation in 60% of *VPS53*_*S790A*_ cells. In contrast, only 30% of *VPS53* and *VPS53*_*S790D*_ expressing cells showed this phenotype (Fig. 5a, black bars). Similar results were obtained for *Vps53*_*S790A*_ expressing cells that were glucose starved (suppl. Fig. 2). Together, these results show that mitochondrial dynamics are changed in cells expressing a non-phosphorylatable mutant of Vps53 grown in the presence of lactate. Co-purification of mitochondria and vacuoles depends on the formation of a contact site between the two organelles (vCLAMP, vacuolar and mitochondrial patch, Elbaz-Alon *et al*., 2014; Hönscher *et al*., 2014). Since mitochondria appeared to be more fragmented in *VPS53*_*S790A*_ cells growing in lactate, we hypothesized that the number of vCLAMPs remained constant under these conditions. To test this, we used the established vCLAMP sensor strain. The vCLAMP split-Venus fluorescent reporter strain contains the N-terminus of the Venus fluorescent protein fused to the transmembrane protein Tom70, a subunit of the TOM complex, and the C-terminus of Venus fused to Zrc1, a zinc channel of the vacuole membrane (González Montoro *et al*., 2018). We first quantified total cellular fluorescence of the reporter in Vps53, *Vps53*_*S790A*_ and *Vps53*_*S790D*_ grown in the presence of glucose. Under these conditions, we did not detect any significant changes in the abundance of vCLAMP signals (Fig. 5b, white bars). Previous studies determined a decrease of vCLAMPs after 26h exposure to respiratory growth conditions (Hönscher *et al*., 2014). In contrast, in cells grown 90 minutes in the presence of lactate we detected a slight increase in vCLAMP signals in cells carrying the *VPS53*_*S790A*_ allele (Fig. 5b, black bars). A constant number of vCLAMPs in lactate growing cells together with a higher degree of mitochondrial fragmentation explains the decreased amount of mitochondrial proteins identified in our SILAC purification of vacuoles from *Vps53*_*S790A*_ cells.

**Figure 5:**
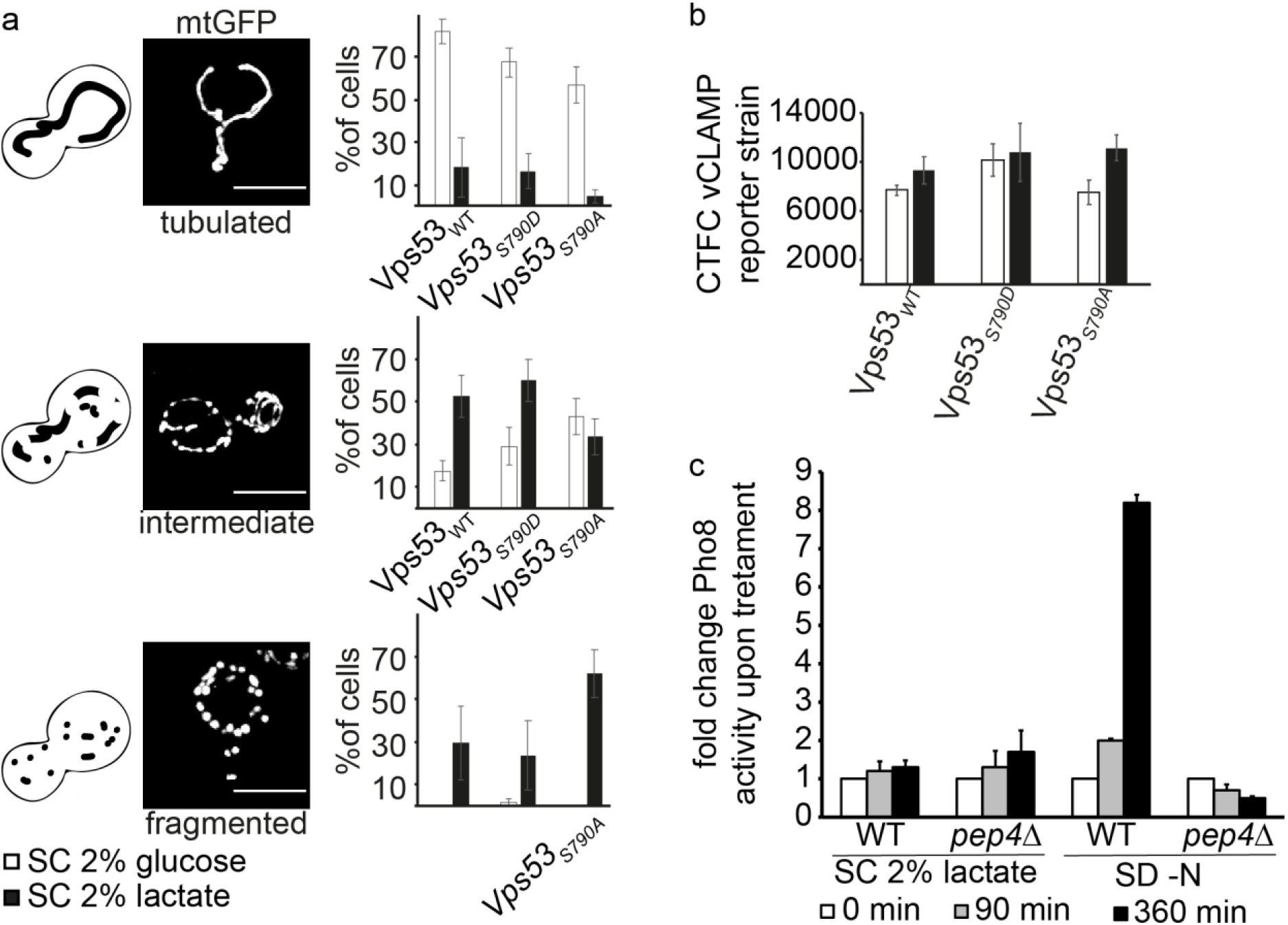
Vps53 phosphorylation causes mitochondrial phenotypes. **(a)** Vps53 phosphorylation affects mitochondrial morphology. Mitochondria were visualized with mtGFP in *vps53Δ* cells expressing *VPS53, VPS53*_*S790A*_ and *VPS53*_*S790D*_ from plasmids. Exponentially growing cells were shifted to media with 2% lactate as a non-fermentable carbon source. Live images were acquired after 90 min (Scale bar 5 μm). Mitochondrial morphology was classified as follows: fragmented, mainly small and spherical; intermediate, mixture spherical and shorter tubulated; and tubulated, interconnective and elongated. Quantification of the indicated mitochondrial morphology was determined as percentage of the total number of counted cells. Cells grown in SDC showed the dominant phenotype of tubulated mitochondria that shifts to intermediate and fragmented structures during respiratory growth condition. *VPS53*_*S790A*_ cells (∼70%) show a higher rate of fragmentation compared to WT and *VPS53*_*S790D*_ cells. *VPS53*_*S790D*_ and WT cells have comparable phenotypes. Approximately 58% of WT and ∼65% *VPS53*_*S790D*_ cells show intermediate morphological structure and slightly increased levels of fragmentation during respiratory growth. Error bars represent the average of three independent experiments (150 ≥ cells), **(b)** A vCLAMP reporter consisting of the VN tagged mitochondrial membrane protein Tom70 and the vacuolar Zrc1 tagged with the complementary split Venus half. *vps53Δ* cells were complemented with plasmids containing either the WT or VPS53_S790A_and *VPS53*_*S790D*_ mutants. VPS53_wt_ was grown to mid-log phase and shifted to lactate containing media for 90 min and imaged. The corrected total cell fluorescence (CTCF) was measured for each cell. CTCF values for respiratory growth conditions (black bars) and control (white bars) are given as means with standard deviation (n = 3). **(c)** Growth on lactate as a sole carbon source does not induce mitophagy. WT and *pep4Δ* strains expressing the mitophagy reporter *mtto-pho8Δ60* were subjected to respiratory growth conditions (SC-0.5% lactate) or nitrogen starvation (SD-N) for the times indicated. The fold change of specific mito-pfio8Δ60 activity relative to glucose-grown control cells is given as mean +/- s.d. (n = 3). Only nitrogen starvation induced mito-phio8Δ*60* activity in a *PEP4*-dependent.

It was previously shown that prolonged growth of yeast cells in lactate can induce mitophagy (Kanki *et al*., 2009). To rule out diminished mitophagy as the cause for decreased co-purification of mitochondria in our vacuole purifications we directly analyzed mitophagy under our growth conditions. Therefore, we generated strains, in which the autoinhibited phosphatase Pho8Δ60 is targeted to mitochondria. When mito-Pho8Δ60 reaches the vacuole as a result of autophagy, its inhibitory pro-peptide is cleaved off in a Pep4-dependent manner and its phosphatase activity is unmasked, allowing quantitative assessment of mitophagy (Campbell and Thorsness, 1998; Schuck *et al*., 2014). We switched yeast cells from glucose-containing medium to lactate-containing medium for up to 6 h to potentially activate mitophagy. However, we could not detect induction of mitophagy (Fig. 5c, left part). In contrast, mito-Pho8Δ60 activity was strongly induced in WT but not in *pep4Δ* cells when autophagy was triggered by nitrogen starvation, demonstrating functionality of the assay (Fig. 5c, right part Kanki *et al*., 2009). Our measurements show that autophagy is not occurring in our culture conditions, excluding the hypothesis that the decreased presence of mitochondrial proteins in *Vps53*_*S790A*_ vacuoles is due to an impaired autophagy.

### Phosphorylation of Vps53 affects Golgi mitochondria proximity

How does Vps53 phosphorylation affect mitochondrial morphology? When we analyzed the co-localization of Vps53 with a Golgi marker (sec7-mKate) and mito-BFP, we observed partial co-localization of these structures in *Vps53*_*S790D*_ expressing cells grown in the presence of glucose (Fig. 6a). Interestingly, shifting Vps53-GFP cells to lactate containing medium also led to close proximity of the Golgi apparatus and mitochondria (Fig. 6b). We therefore asked whether Vps53 could be involved in the formation of a contact site between the Golgi apparatus and mitochondria. To test this, we used an assay based on a fluorescence reporter strain that monitors the extent of organelle proximity by split-Venus complementation (Shai *et al*., 2018). When one half of the split Venus system was fused to Tom70 and the other half to Vps53 we detected robust fluorescent signals with an average of three structures per cell in glucose grown cells. In comparison, *Vps53*_*S790A*_ expressing cells also showed signals, but the amount of cells without a signal was higher. V*ps53*_*S790D*_ expressing cells showed higher signal with an average of seven structures per cell and no cells without any signal (Fig. 6c and d). When we switched cells to lactate containing medium for 90 minutes, we observed an increase of fluorescent signal in WT cells. The signal in both phospho-mutants of Vps53 remained constant, suggesting that the formation of the contact site is indeed depending on the phosphorylation status of Vps53 (Fig. 6e and f). If the formation of a contact site between the Golgi apparatus and mitochondria is dependent on the phosphorylation of Vps53, we expected to detect proximity of the Golgi apparatus and mitochondria based on a different set of organelle markers. We therefore fused the C-terminal half of split Venus to the trans-Golgi marker Sec7 and the N-terminal half to the mitochondrial marker Tom70. In cells expressing a WT copy of Vps53 we detected a signal in about 75% of the cells with an average of two structures per cell. When *Vps53*_*S790A*_ was expressed we failed to detect any signal (Fig. 6g and h). In contrast, *Vps53*_*S790D*_ expressing cells had on average four fluorescent structures and signal was detectable in 100% of the cells. Switching cells to lactate containing medium increased signal levels in WT cells, while the signal in both phospho-mutants remained constant (Fig. 6i and j). Interestingly, deletion of the recruiter Rab of the GARP complex, Ypt6, increases the formation of contact sites (suppl. Fig. 3), which most likely is caused by a dispersal of the Golgi complex the *YPT6* knockout (Conibear and Stevens, 2003). Together, our results show that the formation of a contact site between the Golgi apparatus and mitochondria (GoMiCS) depends on the GARP subunit Vps53 and its phosphorylation.

**Figure 6:**
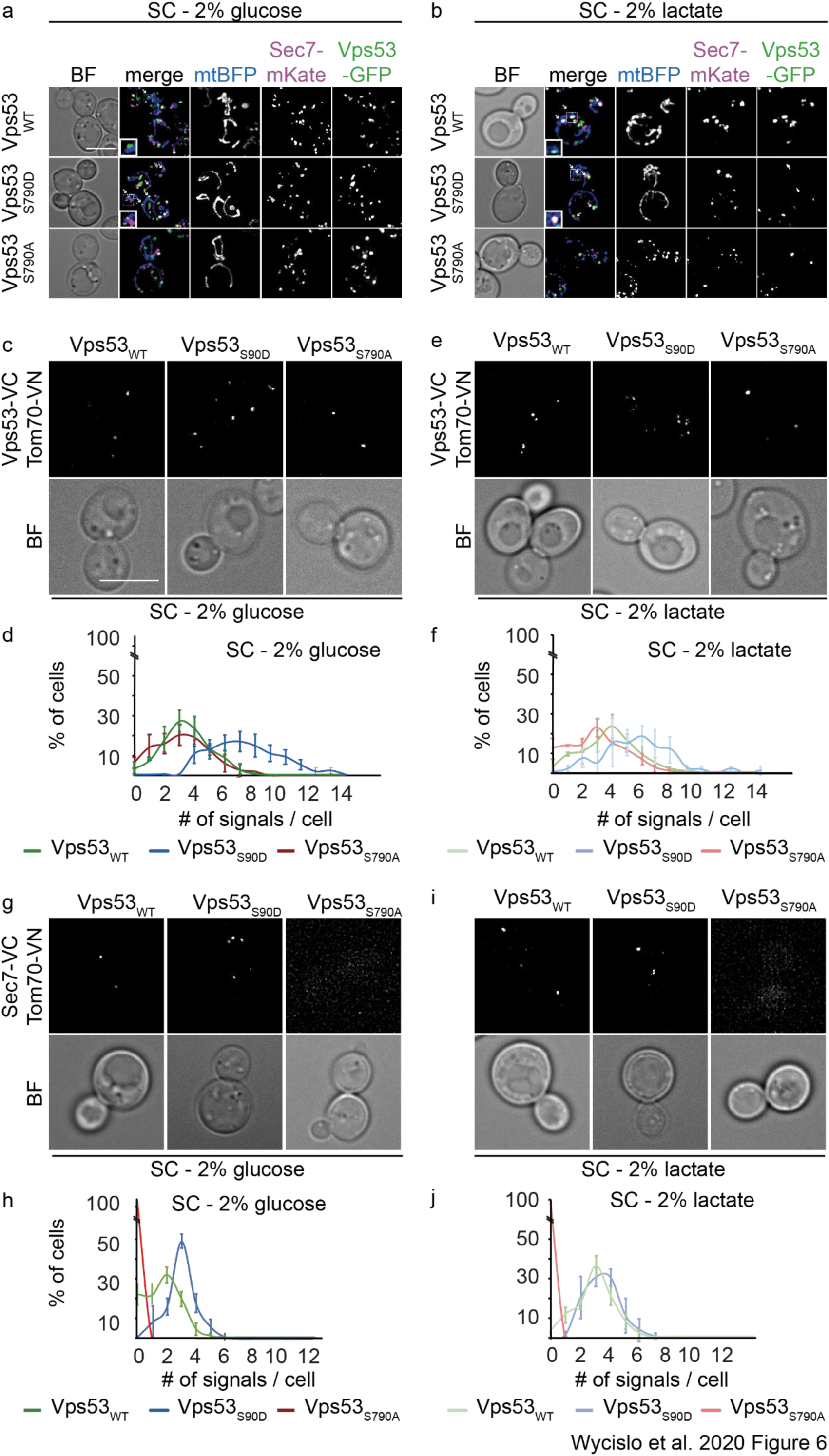
Vps53 controls the formation of a Golgi-Mitochondria contact site. (**a,b**) Vps53-GFP colocalizes with the mitochondrial marker mtBFP and the trans Golgi marker Sec7. Subcellular localization of the GFP tagged Vps53 and phospho-mutant variants (green) was analyzed in cells co-expressing the mitochondrial marker mtBFP (blue) and the mKate tagged trans Golgi marker Sec7 (magenta). Cells were grown to mid-log phase and shifted for 90 min in lactate containing medium and imaged to assess co-localization (Scale bar 5 μm). Colocalization of Sec7-mKate and Vps53-GFP variants was not affected either in SC-D media (**a**) or under respiratory growth (**b**) conditions (SC 2% lac.). Lactate treatment leads to re-localization of Vps53-GFP foci to mitochondria. Arrowheads indicate co-localization of Vps53-GFP variants with mitochondria and Sec7-mKate. Magnification of co-localization (boxed region) are shown on the left corner of the merged image. **(c)** Analysis of the proximity of Vps53 with mitochondria. Cells expressing the mitochondrial membrane protein Tom-70 tagged with one half of the split Venus protein together with either Vps53, *Vps53*_*S790*A_,*Vps53*_*S790D*_ expressing the complementary half of split venus were grown in the presence of glucose. Scale bar = 5μM **(d)** Quantification of the observed spots per cell from **(c). (e)** Cells expressing the mitochondrial membrane protein Tom-70 tagged with one half of the split Venus protein together with either Vps53_WT_, *Vps53*_*S790*A_, *Vps53*_*S790D*_ were grown for 90 min in the presence of lactate. Scale bar = 5μM **(f)** Quantification of the observed spots per cell from (e). The average number of signals per cell increases only in Vps53_WT_ expressing cells and stays constant in *Vps53*_*S790A*_ and *Vps53*_*S790D*_ expressing cells. **(g)** Analysis of the proximity of the Golgi with mitochondria. Cells expressing the mitochondrial membrane protein Tom-70 tagged with one half of the split Venus protein together with the Golgi marker Sec7 expressing the complementary half of split venus were grown in the presence of glucose. Vps53_WT_ (left panel), *Vps53*_*S790*D_ (middle panel), *Vps53*_*S790A*_ (right panel) were co-expressed in the cells. Scale bar = 5μM **(c). h)** Quantification of the observed spots per cell from (g). **(i)** Cells expressing the mitochondrial membrane protein Tom-70 tagged with one half of the split Venus protein together with the Golgi marker Sec7 expressing the complementary half of split venus were grown for 90 min in the presence of lactate. Vps53_WT_ (left panel), *Vps53*_*S790*D_ (middle panel), *Vps53*_*S790A*_ (right panel) were co-expressed in the cells. Scale bar = 5μM. **(j)** Quantification of the observed spots per cell from (i). The average number of signals per cell increases only in Vps53_WT_ expressing cells and stays constant in *Vps53*_*S790A*_ and *Vps53*_*S790D*_ expressing cells.

## Discussion

In this study we identify the Vps53 subunit of the GARP complex as a target of the yeast AMP kinase homolog Snf1. Vps53 is specifically phosphorylated at serine 790 and phosphorylation depends on the availability of a carbon source (low versus high glucose) and the nature of the carbon source (glucose versus lactate). Surprisingly, Vps53 phosphorylation does not affect GARP complex assembly or any trafficking pathway, but instead affects mitochondrial dynamics. We propose that this phenotype is caused by the lack of the formation of a novel, previously unrecognized membrane contact site between the Golgi apparatus and mitochondria, that we name GoMiCS (for <u>Go</u>lgi-<u>Mi</u>tochondria <u>C</u>ontact <u>S</u>ite).

The phosphorylation site on Vps53 is at the very C-terminal end of the protein. Based on the crystal structure of the C-terminal fragment of Vps53, this site is in a disordered region of the protein (Vasan *et al*., 2010). The C-terminal part of Vps53 folds into two adjacent helical bundles, one formed by α-helices 1 to 3 and a second one formed by the α-helices 4 to 9. The identified phosphorylation site on serine 790 is located close to the end of the ninth α-helix at position 778. Based on the cryo-EM structure of the intact GARP complex, the C-terminus of Vps53 is distant from the core where all four subunits interact with each other (Chou *et al*., 2016). Thus, the phosphorylated region of Vps53 appears to be accessible to the Snf1 kinase. Additionally, based on the overall subunit architecture of the complex, it is unlikely that phosphorylation of Vps53 affects the overall formation of GARP. The C-terminal part of Vps53 is also not important for the interaction with any effector proteins of GARP. The recruiter RAB of GARP is Ypt6 and this interaction was previously shown to occur between the Vps52 subunit and Ypt6 (Siniossoglou *et al*., 2000; Siniossoglou and Pelham, 2001). However, deletion of Ypt6 causes dispersion of GARP in the cells but does not diminish its association with a membrane fraction. The interaction with the SNARE protein Tlg1 is mediated by the Vps51 subunit of GARP (Siniossoglou and Pelham, 2001; Conibear and Stevens, 2003).

Based on our results it also appears unlikely that the C-terminal part of Vps53 nor its phosphorylation is important for any known trafficking pathway in yeast. The GARP complex has been shown to be important for CPY transport based on a defect in Vps10 CPY receptor recycling (Conboy and Cyert, 2000; Conibear and Stevens, 2000; Vasan *et al*., 2010; Bonifacino and Hierro, 2011). Our analysis shows that neither the non-phosphorylatable mutant of Vps53 nor the phospho-mimetic mutant of Vps53 affect CPY trafficking. In addition, depletion of the carbon source or growth in a non-fermentable carbon source do not affect CPY trafficking in our studies. We and others have previously shown that the GARP complex is important for the recycling of several plasma membrane proteins, including the SNARE Snc1 and members of the aminophospolipid-flippase family, such as Dnf1 (Siniossoglou and Pelham, 2001; Conibear and Stevens, 2003; Takagi *et al*., 2012; Eising *et al*., 2019). Glucose starvation has been suggested to have a direct effect on the abundance of plasma membrane proteins by downregulation of the recycling pathway (Lang *et al*., 2014). Our results suggest that this pathway is independent of Vps53 phosphorylation.

Instead of any trafficking-related phenotypes, we find that Vps53 phosphorylation affects mitochondrial morphology. In a non-phosphorylatable Vps53 mutant, mitochondria fragmented significantly faster compared to cells expressing a WT version or a phospho-mimetic version of Vps53. Fragmentation of fission yeast mitochondria upon glucose depletion has been shown before (Zheng *et al*., 2019). However, long term exposure to respiratory growth conditions result in a tabulated mitochondrial network (Egner *et al*., 2002). The opposite effects we observe after 90 minutes of growth in lactate could be part of a remodeling process to adapt to different bioenergetics (Westermann, 2012). At the moment we can only speculate about the role of Vps53 and phosphorylation by Snf1 in this process. Our data suggest that phosphorylation of Vps53 leads to the formation of a membrane contact site between the Golgi-apparatus and mitochondria, which we coined GoMiCS. While membrane contact sites have been observed between almost all organelles in the cell (Shai *et al*., 2018), GoMiCS have been undetected so far. One reason could be the highly dynamic behavior of both the Golgi apparatus and mitochondria in yeast. Similar to other studies, we were only able to identify the described contact site by a microscopy-based fluorescence complementation assay. However, it is important to emphasize that *i)* we observe GoMiCS with proteins expressed under their endogenous promotor and *ii)* the exchange of a single amino acid in Vps53 leads to the loss of the contact site. These findings argue against an artificial induction of the GoMiCS by the tags used for the complementation assay.

Our observation of a Vps53 role in formation of GoMiCS agrees with previous findings on additional functions of subunits in tethering complexes. As another example, the Vps39 subunit of the vacuolar HOPS tethering complex also mediates the formation of a contact site, here between the yeast vacuole and mitochondria (Elbaz-Alon *et al*., 2014; Hönscher *et al*., 2014). In addition, the formation of this contact site is dependent on the available carbon source and Snf1, where respiratory growth conditions led to the reduction of the contact site after long term exposure to respiratory growth conditions. The physiological consequences of this reorganization of membrane contact sites, for example regarding the transfer of lipids between organelles, remain to be explored.

Membrane contact sites have been described as regions of vesicle-independent transport of lipids or other metabolites between adjacent organelles. We originally identified Vps53 phosphorylation to be decreased in sphingolipid depleted cells (Fröhlich *et al*., 2016). Does formation of GoMiCS play a role in sphingolipid transfer between the Golgi and mitochondria and thus affect sphingolipid homeostasis? AMPK activation has been linked to sphingolipid homeostasis but mechanistic insights are lacking (Liu *et al*., 2013). Our data suggest that Snf1 activation upon carbon source depletion could be attenuated by additional inhibition of sphingolipid metabolism by myriocin. However, the signaling pathway regulating Snf1 upon sphingolipid depletion remains elusive. In addition, the GARP complex plays an important role in maintaining sphingolipid homeostasis (Fröhlich *et al*., 2015; Petit *et al*., 2020). In our current study we do not observe an influence of Vps53 phosphorylation on sphingolipid metabolism. However, the only sphingolipid modifying enzyme located at mitochondria is Isc1, the yeast homolog of a neutral sphingomyelinase (Kitagaki *et al*., 2007). The exact function of Isc1 at mitochondria remains largely elusive, since complex sphingolipids, the substrate of Isc1, are not found there. But Isc1 has been implicated in mitochondrial fission and mitophagy (Teixeira *et al*., 2015).

In contrast to sphingolipids Phospshphatidylethanolamine (PE) is generated in mitochondria by the phosphatidylserine decarboxylase Psd1. The required phosphatidylserine is likely be transported via membrane contact sites towards mitochondria (Aaltonen *et al*., 2016). In addition, Psd1 generated PE has been implicated in mitochondrial fusion (Chan and McQuibban, 2012; Joshi *et al*., 2012). If the mitochondrial remodeling we observe after 90 min of lactate exposure is the starting point of mitochondrial fusion, additional PE import could be required (Calzada *et al*., 2019). Vps53 dependent GoMiCS could therefore be an additional source for of PE import into mitochondria. Along that line, the second phosphatidylserine decarboxylase Psd2 is localized at the Golgi/endosomal system (Kitamura *et al*., 2002; Gulshan *et al*., 2010).

Since the C-terminal part of Vps53 that carries the identified phosphorylation is not conserved in mammalian cells, it is unclear if the mammalian GARP complex has a similar function in the formation of a membrane contact site. Interestingly, organelle contacts between the Golgi and mitochondria have been described (Dolman *et al*., 2005; Valm *et al*., 2017) and a role in calcium signaling has been suggested. Whether formation of these contacts is GARP dependent remains to be studied in the future.

## Materials and Methods

### Yeast strains and plasmids

Yeast strains used in this study are described in supplementary file 1. Plasmids used in this study are summarized in supplementary file 2. Genetic manipulations were made by homologous recombination of PCR fragments as described previously. For the generation of endogenously expressed GFP tagged Vps53 phospho mutants, WT strains were first transformed with the tag and a resistance marker according to standard procedures. To introduce the genomically encoded phospho mutants the transformed yeast strain was used to amplify parts of the Vps53 c-terminus together with the GFP tag and the resistance marker. Mutations were introduced with the forward primer. The resulting PCR products were used to transform WT strains. Introduction of the tag and resistance marker were confirmed by PCR. Introduction of the mutation was checked vie PCR amplification and sequencing.

### Yeast media and growth conditions

Yeast cells were grown in rich media YP + X containing the respective carbon source 2% (w/v) D-glucose (X = D) or 2% (w/v) galactose (X = G); 1% (w/v) yeast extract and 2% (w/v) Peptone or in selective synthetic minimal media (SD + Y; 0.67% yeast nitrogen base without amino acids, 2% carbon source and supplemented amino acids).

Lactic acidic stress was induced by shifting cells (OD_600_ 0.8) to YP + or SC + X media containing 2% lactate or 0.5% lactate. Glucose starvation was induced by shifting cells to medium containing 0.05% glucose. The respective percentage (w/v) of carbon source was used and cells were incubated for 90 min at 30 °C unless otherwise noted.

For SILAC labeling procedures, lysine auxotroph strains were grown in SC-D-lysine medium consisting of 2% glucose or 2% lactate, 6.7 g/L yeast nitrogen base without amino acids and yeast synthetic dropout without lysine (Sigma Aldrich). Pre-cultures were grown over night in the presence of 30 mg/L normal lysine or heavy lysine (L-Lysine ^13^C_6_^15^N_2_; Cambridge Isotope Laboratories) and diluted to OD_600_ 0.1. Cells were grown to OD_600_ 0.5-1 before harvest.

### Fluorescence Microscopy

Cells were grown to logarithmic phase in synthetic selective media (SC-D) containing 2% glucose supplemented with essential amino acids. Where indicated, cells were switched to respiratory growth conditions (X = 2% lac) and starvation (X = 0.002% glc) was induced by transferring them at OD_600_ 0.8 to SC-X medium for 90 min at 30 °C unless otherwise noted. Cells were imaged live in SC medium unless stated otherwise on an Olympus IX-71 inverted microscope equipped with 100 x NA 1.49 and 60x NA 1.40 objectives, a sCMOS camera (PCO, Kelheim, Germany), an InsightSSI illumination system, BF (bright field); 4′,6-diamidino-2-phenylindole; GFP; mCherry and Cy5 filters, and SoftWoRx software (Applied Precision, Issaquah, WA). We used z-stacks with 350 nm spacing for constrained-iterative deconvolution (SoftWoRx). All microscopy image processing and quantification was performed using ImageJ (National Institutes of Health, Bethesda, MD; RRID:SCR_003070). Maximum intensity Z projections are shown unless otherwise noted. Where indicated, at least 50 cells from 3 independent experiments were quantified for the relevant statistics.

Visualization organelle proximity using bimolecular fluorescent complementation (BifC) assay. For juxtaposition Vps53 or late Golgi-mitochondria 3D image stacks of 6 images were collected with 350 nm z-increment. Positive signals within the cell volume shown in (Fig. 6) were determined using a manual and automatic method. Manual method: minimum thresholds were set manually and the gausssian blur with a sigma radius of 0.5 was applied to minimize the cytosolic signals. For the automated method, the quantification of foci was performed in ImageJ (national Institutes of Health, Bethesda, MD) by a self-written image processing application combining built-in routines and the “3D Objects counter” plug-in. Briefly, single cells were segmented from the bright field image and Venus signals within the deconvolved image stack were counted based on the plug in “3D Objects Counter”. For the optimization of the signal-to-noise ratio was each signal within the stack convolved with the standard deviation sigma of a Gaussian function (Gaussian-Blur), whereby sigma was given in units corresponding to the pixel scale, in addition background signals were removed by selecting a background-area. Signal intensities were normalized and convolved with the mean of the Venus kernel. Foci segmentation within the stack was restricted to a maximum threshold of 180 and to a minimum voxel value of 10. Objects were displayed and counted in a map of all z-planes. Maximum intensity z projections of z planes with signals are shown. Raw images were quantified and are shown. Graphs show mean ± SD for three independent experiments. vCLAMP reporter strain images shown are deconvolved images of one z-plane. For the quantification of the signal in the reporter strain. The total cell fluorescence per cell of the sum intensity z projection was quantified with ImageJ. The formula CTCF = Integrated Density – (Area of selected cell X Mean fluorescence of background readings) was used to calculate the corrected total cell fluorescence. Graphs show mean ± SD for three independent experiments. Mitochondria morphology was examined by 3D projection. Deconvolved z-stacks of 12 images with 350 nm spacing were used. Cells bearing plasmid pXY142-mtGFP were grown in selective synthetic medium without Leucin (Leu). Graphs show mean ± SD for three independent experiments.

### Mitophagy assay

Strains were grown to mid-log phase in glucose-containing YPD medium. Ten OD_600_ units each were collected as control samples. Cells were washed once with water and resuspended to an OD_600_ of 0.5 in SC + 0.5% lactate or nitrogen-free SD-N medium. The *pep4Δ* strain was additionally treated with 1 mM PMSF to completely block vacuolar proteolysis. Ten OD_600_ units each were harvested after 90 and 360 min. The Pho8 assay was performed as described (Schäfer and Schuck, 2020) with lysates adjusted to 4 ODs/100 μl and a reaction time of 25 min. Specific phosphatase activities were normalized to the activities in the corresponding control samples to obtain fold changes.

### Liquid CPY-Invertase assay

Invertase deficient W303 *suc2Δ* cells were grown according to the standard procedures in YP 2% fructose medium. Cells were shifted for 90 min to YP 2% lactate containing medium. 10 OD_600_ units were used for cells expressing Vps53, *Vps53*_*S790A*_ and *Vps53*_*S790D*_. For strains deleted for endogenous Vps53 expressing empty pRS306 vector 0.5 OD_600_ units were used. The assay was performed according to the procedure used by (Dalton *et al*., 2015). Results are reported as units of activity per 1 OD_600_ for cells grown under respiratory growth conditions and control cells. Error bars represent the standard deviation for three biological experiments, and three technical replicates were analyzed. Comparison between cells expressing the WT form and the phospho-mutants Vps53 were made using unrelated t-test. A p value <0.005 was consider significant.

### *In vitro* kinase assays

Recombinant GST-Vps53_552-811_ and active Elm1^KD^ were incubated with recombinant wild type Snf1^KD^ or the kinase dead mutant (T210A). Kinases and substrate were mixed at a 1:1:5 ratio in Phosphorylation-Buffer [10 mM Tris (pH 7.5), 10 mM MgCl2, 0.4 mM EDTA, 10% glycerol, 1 mM ATP] and incubated at 30 °C for 30 min and gentle shaking (300 – 500 rpm). For mass spectrometry analysis 50 μl of denaturation buffer was added.

### Protein Expression and Purification

Recombinant GST tagged Vps53 C-terminal fragment (residue 552-822) was purified for the kinase assay from 1 L [LB, 100 μg/ml Ampicillin and 100 μg/ml Chloramphenicol] *E. coli* BL21 (DE3) Rosetta cells after induction [OD_600_ 0.8] by 0.5 IPTG (isopropyl β-D-thiogalactosidase). Cells were cultured for 16 h at 16 °C. Cell lysis and extraction were carried out at 4 °C on ice. Cells were harvested [5000 rpm, 10 min] and the pellet was washed once with 10 pellet volume Wash-Buffer (WB) [PBS (pH 7.0), 1 mM DTT, 0.1% (v/v) NP-40 and 300 mM NaCl] and resuspended in Lysis-Buffer [WB, 0.05 x PIC, 1 mM PMSF]. The cells were lysed, by a Microfluidizer Processor M-110L (Microfluidics) and the lysate was cleared by centrifugation [15 min, 15000 rpm at 4 °C]. Vps53 protein was purified by batch binding to glutathione (GSH)-Sepharose fast flow (GE-Helthcare) [1 h at 4 °C]. The resin was washed five times with 100 bead volume WB [1000 rpm at 4 °C]. To elute immobilized GST-Vps53 recombinant protein, resin was incubated twice with 15 mM and 20 mM reduced GSH for 15 min at 4 °C. Fractions of purification steps and eluate were resuspended in sample buffer boiled for 5 min at 95 °C. Proteins were finally resolved by SDS-PAGE.

The following recombinant proteins were purified according the same procedure as described before. Deviation from the protocol are indicated. The GST-Snf1 kinase domains (wild-type and T210A; residues 1-392) were expressed and purified. The purified protein was eluted with PreScission protease (final volume of 1 ml), incubated at 16 °C for 2 h on a rotating wheel. GST-Vps53 and Snf1 kinase domains eluates were dialyzed against the kinase assay buffer [20 mM Hepes (pH 7.0), 0.5 mM DTT, 5 mM MgOAc]. Glycerol was added to a final concentration of 5% (v/v), and aliquots were stored at −80 °C.

Yeast cells expressing TAP tagged Elm1 kinase domain (residues 421-640) were cultured in 2 L YPG to OD_600_ 3-5 and harvested. The pellet was resuspended in TAP-Buffer [Wash-Buffer, 0.05 x Pic, 1 mM PMSF, 1 mM TCEP] followed by lysis with glass beads in a FastPrepTM machine (MP Biomedicals). Lysate was centrifuged for 20 min, 4000 rpm at 4 °C followed by ultracentrifugation [35000 rpm, 1.10 h]. Supernatant was incubated with IgG Sepharose 6 Fast Flow (GE Healthcare) for 2 h at 4 °C on a nutator. IgG resin was washed with 15 mL Wash-Buffer [PBS (pH 7.0), 10% (v/v) glycerol, 20 mM Chaps, 1 mM NaF, 1 mM Na3VO4, 20 mM β-Glycerophosphate]. TEV cleavage of immobilized protein was performed in 400 μL Cleavage-Buffer [PBS (pH 7.0), 10% Glycerol, 20 mM Chaps] over night at 4 °C on a turning wheel. Detergent from the eluate was removed by Detergent Removal Spin Column (Pierce).

### Vacuolar isolations

Respiratory growth conditions were induced at OD_600_ 0.8 for 90 min. Vacuoles were purified from 2 L. Same OD-units of SILAC labeled cell cultures were mixed and harvested together and the pellet treated with Tris-buffer (0.1 M Tris, pH 9.4; 10 mM DTT) and spheroplasting buffer (0.6 M sorbitol, 50 mM KPi, pH 7.4, in 0.2x YPD). After lyticase digestion, vacuoles were isolated via dextran lysis and Ficoll gradient flotation (Cabrera and Ungermann, 2009). 500 μl of the 0-4% interphase were taken and mixed with 25 μl 20x PIC. Protein concentration was measured by a Bradford assay.

### GFP pulldown assays

Cells were grown in the presence of “heavy” lysine (l-lysine-U-13C6,15N2) and WT cells were grown in the presence of normal, “light” lysine. We harvested 500 OD600 units of cells by centrifugation and resuspended them in 500 μl of lysis buffer (150 mM KOAc,20 mM 4-(2-hydroxyethyl)-1-piperazineethanesulfonic acid [HEPES], pH 7.4, 10% glycerol, and complete protease inhibitor cocktail [Roche, Basel, Switzerland]). Zirconia beads (500 μl, 0.1-mm diameter; BioSpec Products, Bartlesville, OK) were added and cells were lysed using a FASTPREP (MP Biomedicals, Solon, OH) for 60 s at 4°C. Beads were removed by centrifugation, and Triton X-100 was added to a final concentration of 1%. After a 30-min incubation at 4°C, lysates were cleared by centrifugation for 10 min at 1000 × *g*. Equivalent amounts of “light”-labeled control and “heavy”-labeled *GFP–*containing lysates or “light”-labeled control and “heavy”-labeled *GFP–*containing lysates were incubated (separately) with GFP-Trap agarose beads (Allele Biotechnology, San Diego, CA) for 30 min at 4°C. Beads were washed three times with lysis buffer and three times with wash buffer (150 mM NaCl, 20 mM HEPES, pH 7.4). Beads from *GFP* pull downs and control pull downs were combined in 100 μl of denaturation buffer.

### Proteomics

Mass spectrometry was done with purified vacuoles and whole cell lysate. Vacuoles were further purified by in-gel digest and cell lysate samples by Filter Aided Sample Preparation (FASP; Wiśniewski *et al*., 2009). Purified vacuole samples were precipitated with 100% TCA and the protein pellet washed with acetone. The pellet was solved in 4x loading dye and loaded on a 10% denaturating SDS-gel for some minutes.

For in gel digestion, gel pieces with proteins were excised and incubated in destaining buffer [25 mM NH4HCO3 (ABC) / 50% EtOH] twice for 20 min at 25°C under shaking. After dehydration in 100% EtOH [twice for 10 min at 25°C] and drying the gel pieces were rehydrated in reduction buffer [10 mM DTT in 50 mM ABC] for 60 min at 56°C followed by alkylation [55 mM iodoacetamide in 50 mM ABC] for 45 min at 25°C in the dark and another washing step for 20 min with digestion buffer. After dehydration in EtOH [10 min, 25°C] and washing with digestion buffer [50 mM NH_4_HCO_3_ in water, pH 8.0, 20 min, 25°C] gel pieces were again incubated twice with EtOH for 10 min and dried. Gel pieces were rehydrated in LysC solution [final 16 μg/ml in 50 mM ABC] for 20 min at 4°C, the excess of solution was removed, digestion buffer added and the sample incubated over night at 37°C. Digestion was stopped by adding 2μl 100% TFA. Gel pieces were incubated twice in extraction buffer [3% TFA / 30% ACN] for 10 min at 25°C and twice with ACN for 10 min at 25°C. The supernatants were collected and dried until most of the solvent was gone and resolved in 50 μl HPLC-grade water.

Cell lysate pellets were lysed in 200 μl lysis buffer [Tris 0.1 M, pH 9; 0.1 M DTT; 5% SDS] for 30 min at 55°C and mixed with 1.2 ml 8 M urea in 0.1 M Tris/HCl pH 8.5 (UA). The cell lysate was centrifuged in a wet filter unit (30.000K) for 15 min at 14.000 rpm and the filter washed four times with 200 μl UA for each 10 min. 200 μl IAA solution [0.05 M iodoacetamide in UA] was added to the filter units, shaken vigorously for 1 min and then incubated for 20 min without mixing in the dark. Samples were washed four times with UA for 10 min at 14.000 rpm and washed again with 50 mM ABC and three times with 20 mM.

For mass spectrometry analysis of GFP-Pulldown, *in vitro* kinase assay and GST-Pulldown samples, in solution digestion was performed. Denaturation buffer was added to the sample (8 M urea, 50 mM Tris-HCl, pH 8, 1 mM dithiothreitol) and incubated for 30 min. Proteins were alkylated by the addition of 5.5 mM iodoacetamide for 20 min in the dark and digested with the endoproteinase LysC or Trypsine overnight at 37°C. The resulting peptide mixture was removed from the beads and desalted following the protocol for StageTip purification (Rappsilber *et al*., 2003).

Reversed-phase chromatography was performed on a Thermo Ultimate 3000 RSLCnano system connected to a Q Exactive*Plus* mass spectrometer (Thermo) through a nano-electrospray ion source. Peptides were separated on 50 cm PepMap® C18 easy spray columns (Thermo) with an inner diameter of 75 μm. The column temperature was kept at 40 °C. Peptides were eluted from the column with a linear gradient of acetonitrile from 10%–35% in 0.1% formic acid for 118 min at a constant flow rate of 300 nl/min. Eluted peptides from the column were directly electrosprayed into the mass spectrometer. Mass spectra were acquired on the Q Exactive*Plus* in a data-dependent mode to automatically switch between full scan MS and up to ten data-dependent MS/MS scans. The maximum injection time for full scans was 50 ms, with a target value of 3,000,000 at a resolution of 70,000 at m/z = 200. The ten most intense multiply charged ions (z=2) from the survey scan were selected with an isolation width of 1.6 Th and fragment with higher energy collision dissociation (Olsen *et al*., 2007) with normalized collision energies of 27. Target values for MS/MS were set at 100,000 with a maximum injection time of 80 ms at a resolution of 17,500 at m/z = 200. To avoid repetitive sequencing, the dynamic exclusion of sequenced peptides was set at 30 s. The resulting MS and MS/MS spectra were analyzed using MaxQuant (version 1.6.0.13, www.maxquant.org/; (Cox and Mann, 2008; Cox *et al*., 2011) as described previously (Fröhlich *et al*., 2013). All calculations and plots were performed with the R software package (www.r-project.org/; RRID:SCR_001905)

### Antibodies

The primary antibodies used in this study were as follows: mouse anti-HA antibody (Roche Life Science), mouse anti-PGK1 monoclonal antibody (Thermo Fisher/Invitrogen), polyhistidine antibody H1029 (SigmaAldrich) to detect Snf1 and phospho-Thr172–AMPK antibody (Cell Signalling Technology). Secondary antibodies were HRP-conjugated (horseradish peroxidase) anti-mouse and anti-rabbit IgG (Santa Cruz Biotechnology, Dallas, TX, United States)

## Acknowledgements

We thank members of the Fröhlich lab, Christian Ungermann and Sebastien Leon for critical comments on the manuscript. Florian Fröhlich is supported by the DFG grant FR 3647/2-2 and the SFB944 (P20). Sebastian Schuck is supported by DFG grant EXC 81.

## Supplementary material

**Supplementary Table 1:** List of all proteins identified in GFP pulldown assays of Vps53-GFP, *Vps53*_*S790*A_-GFP and *Vps53*_*S790D*_-GFP expressing cells.

**Supplementary Table 2:** List of all proteins identified in cell lysates and vacuolar purifications of WT and *Vps53*_*S790A*_ cells grown in the presence of lactate.

**Supplementary Figure 1:**
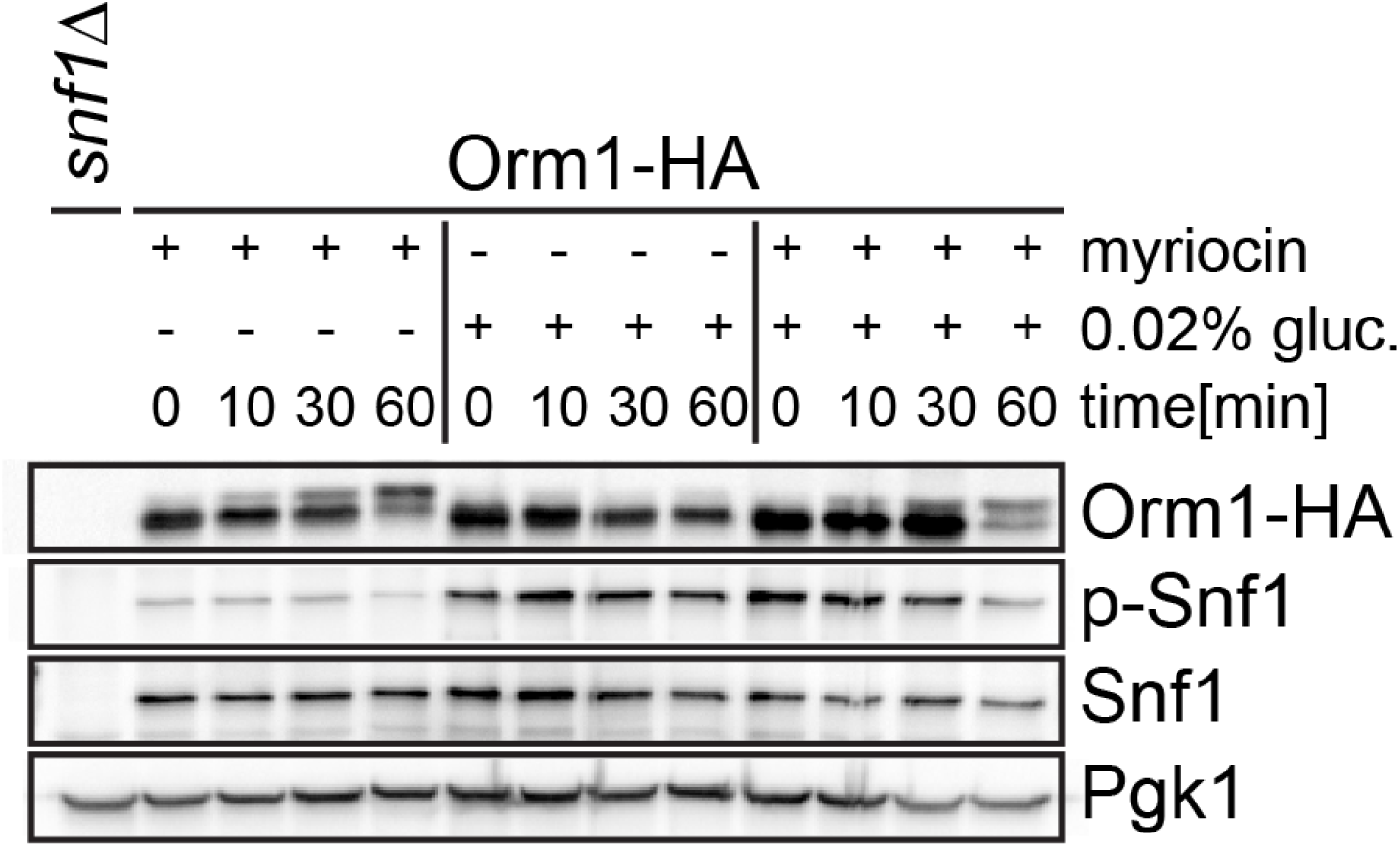
Sphingolipid depletion does not affect Snf1 phosphorylation. Western blot analysis Yeast cells expressing Orm1-HA. Cells were grown in glucose and were untreated (control), depleted from glucose for 60 mins, treated with myriocin for 60 mins or a combination of glucose depletion and myriocin treatment. Samples were collected after 0 mins, 10 mins, 30 mins and 60 mins and analyzed by western blot. Differences in Orm1-HA migration reflect changes in its phosphorylation.

**Supplementary Figure 2:**
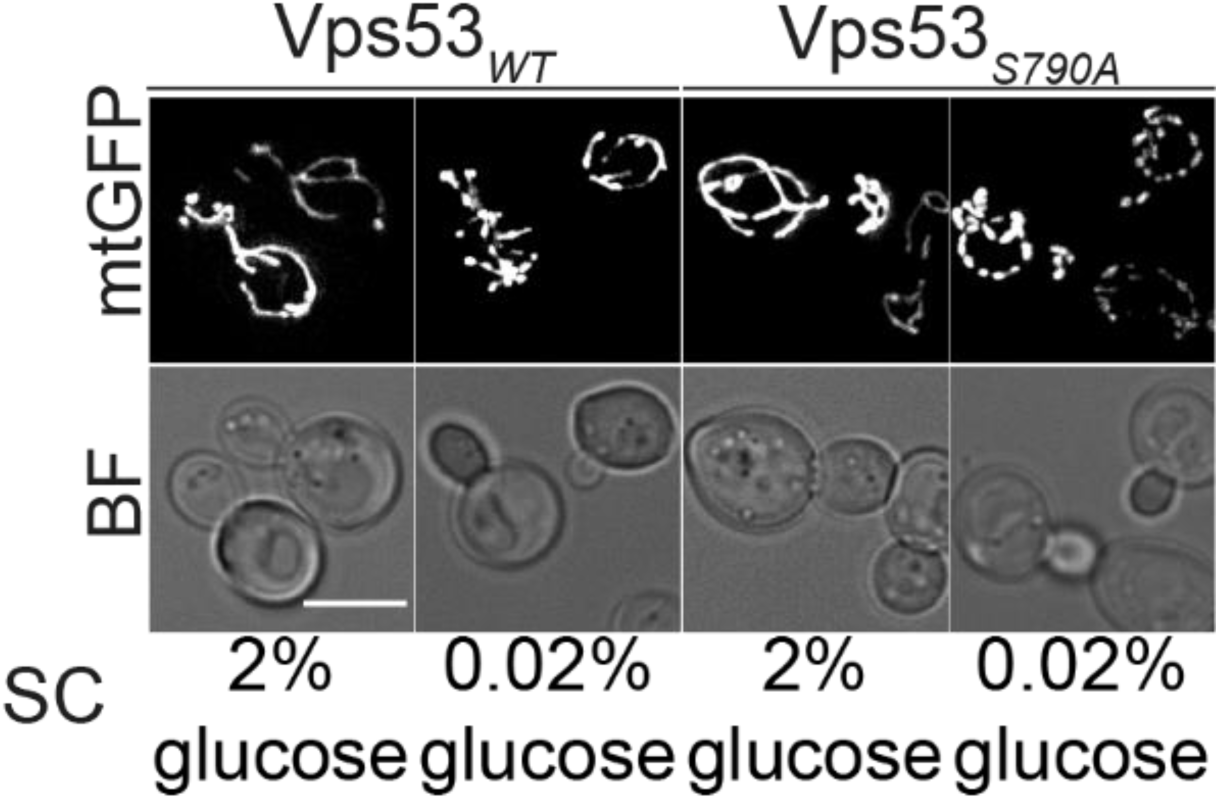
Vps53 mutation unbalances mitochondrial dynamics in glucose starved cells. Mitochondrial morphology was assayed in glucose starved *VPS53* depleted cells co-expressing either Vps53 or *Vps53*_*S790A*_ and a mitochondrial matrix targeted GFP (mtGFP) from a plasmid. Starvation was induced for 90 min in logarithmically grown cells. Glucose starved *Vps53*_*S790A*_ cells displayed fragmented mitochondria.

**Supplementary Figure 3:**
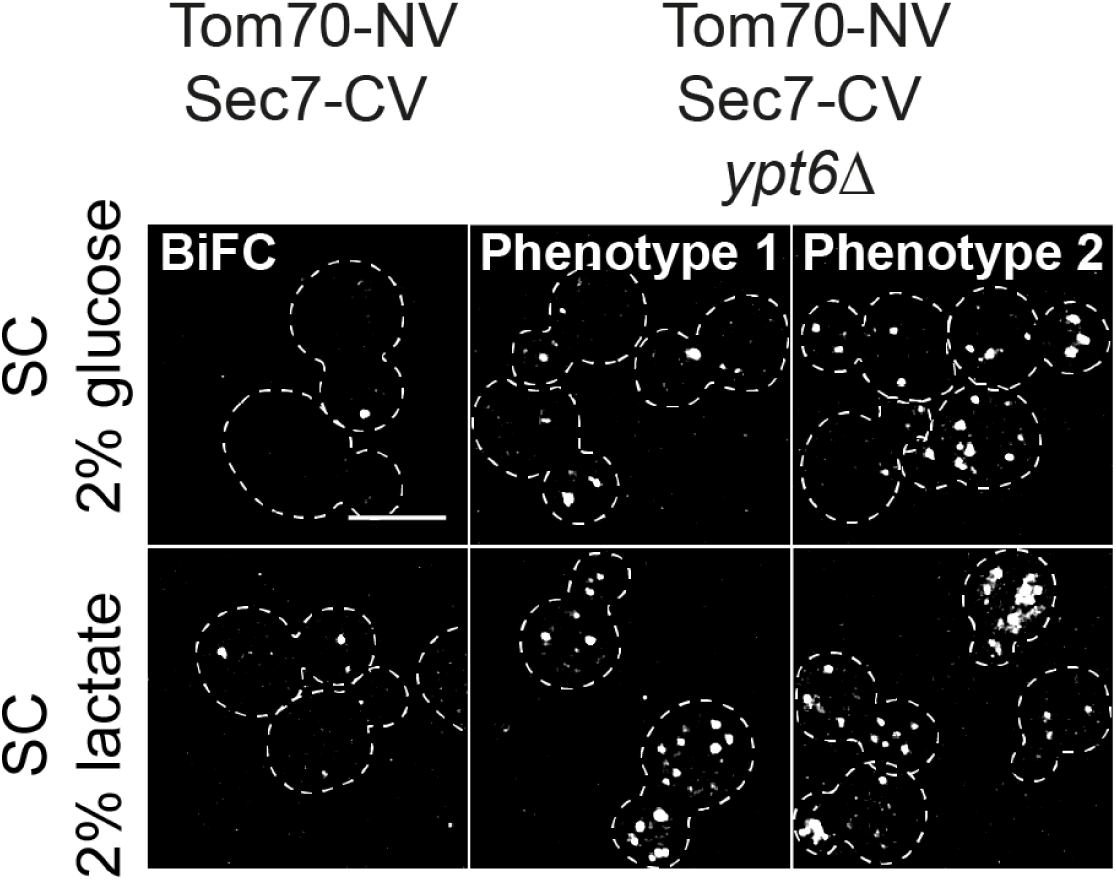
Depletion of the GARP recruiter Ypt6 leads to a considerable increased Venus signal-based GoMiCS formation. Fluorescent complementation based analysis of *ypt6Δ* cells expressing Tom70-NV and Sec7-CV. Logarithmically grown cells were transferred for 90 min to lactate containing medium. The Golgi and mitochondria can be juxtaposed. Loss of the small GTPases Ypt6 affects proximity of the Golgi and mitochondria. Glucose grown cells displayed comparable signals of Venus foci to respiratory Ypt6 expressing cells (phenotype1) and even higher signals (phenotype2). Respiratory growth conditions resulted in considerably more foci (phenotype1) and patch like clusters of complemented Venus signals (phenotype2). Scale bar = 5μM

